# Cell type identification in spatial transcriptomics data can be improved by leveraging cell-type-informative paired tissue images using a Bayesian probabilistic model

**DOI:** 10.1101/2021.11.10.468082

**Authors:** Asif Zubair, Richard H. Chapple, Sivaraman Natarajan, William C. Wright, Min Pan, Hyeong-Min Lee, Heather Tillman, John Easton, Paul Geeleher

## Abstract

Spatial transcriptomics technologies have recently emerged as a powerful tool for measuring spatially resolved gene expression directly in tissues sections, revealing cell types and their dysfunction in unprecedented detail. However, spatial transcriptomics technologies are limited in their ability to separate transcriptionally similar cell types and can suffer further difficulties identifying cell types in slide regions where transcript capture is low. Here, we describe a conceptually novel methodology that can computationally integrate spatial transcriptomics data with cell-type-informative paired tissue images, obtained from, for example, the reverse side of the same tissue section, to improve inferences of tissue cell type composition in spatial transcriptomics data. The underlying statistical approach is generalizable to any spatial transcriptomics protocol where informative paired tissue images can be obtained. We demonstrate a use case leveraging cell-type-specific immunofluorescence markers obtained on mouse brain tissue sections and a use case for leveraging the output of AI annotated H&E tissue images, which we used to markedly improve the identification of clinically relevant immune cell infiltration in breast cancer tissue. Thus, combining spatial transcriptomics data with paired tissue images has the potential to improve the identification of cell types and hence to improve the applications of spatial transcriptomics that rely on accurate cell type identification.

## INTRODUCTION

In the last five years, sequencing-based spatial transcriptomics technologies (1-5) have emerged as a powerful tool to measure spatially resolved genome-wide gene expression directly within tissue sections, offering the potential to interrogate tissue biology in unprecedented detail (6,7). Novel computational methods have already begun to address several analytical challenges posed by these new data, with specific tools developed to identify spatially varying genes (8,9), spatial gene expression patterns (10,11), and cell-cell interactions (12,13). However, the most fundamental problem posed by spatial transcriptomics data—upon which almost all other applications of the data depend—is that of identifying the location and abundance of different cell types (herein referred to as “cell type decomposition”). Several methods have already been developed for this task and generally function by leveraging the expression of a set of cell type-specific marker genes to infer the abundance of each cell type at each slide region (14-17). However, due to statistical multicollinearity (18), cell type decomposition in spatial transcriptomics data will always struggle to differentiate between cell types that are transcriptionally similar. Additionally, low transcript capture, either across an entire slide or in specific regions, can completely impede accurate cell type decomposition.

Here, we present a conceptually novel computational methodology termed Guiding-Image Spatial Transcriptomics (GIST). This method improves cell type decomposition in spatial transcriptomics data by jointly leveraging gene expression data obtained from the spatial transcriptomics platform with cell-type-informative images, for example, AI annotated tissue images, or immunofluorescence stains for cell-type-specific marker proteins, which can be collected, for example, from the reverse side of a tissue section affixed to a spatial transcriptomics capture slide. In a particularly interesting use case, we applied the method to integrate spatial transcriptomics data with deep learning-derived immune cell type annotations in breast cancer pathology slides, where we identified clinically relevant immune cell infiltration that was missed by an initial pathologist’s manual annotation. However, the methodology is generalizable to any spatial transcriptomics platform where informative image-derived cell-type compositional estimates can be obtained. Thus, combining spatial transcriptomics and paired tissue images has the potential to improve all applications of spatial transcriptomics data that rely on the accurate annotation of cell types, such as estimating cell type specific differential expression (19).

## MATERIALS AND METHODS

### Technical details of the GIST statistical model

The expression of gene *i* at each spatial transcriptomics mRNA capture spot *j* is assumed to be approximately a weighted sum of the average expression of that gene in each of the cell types captured by that spot. If our spatial transcriptomics data are arranged in a matrix ***Y***, where the rows represent *i* = 1, …, *m* genes and the columns represent *j* = 1, …, *n* spots, then this relationship can be summarized by the following equation (Fig. 1):

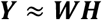

where ***W*** is an *m* × *p* matrix of cell type specific gene expression signatures, approximating the average expression of each gene in each cell type in this tissue, with each column of ***W*** representing one of the *pp* cell types and each row representing one of the *mm* genes. ***W*** is a *p* × *n* matrix of cell type proportions (or probabilities if the data are subcellular resolution) where each column ***W***^(*j*)^ represents the proportions of each of *p* cell types at spot *j*.

**Figure 1:**
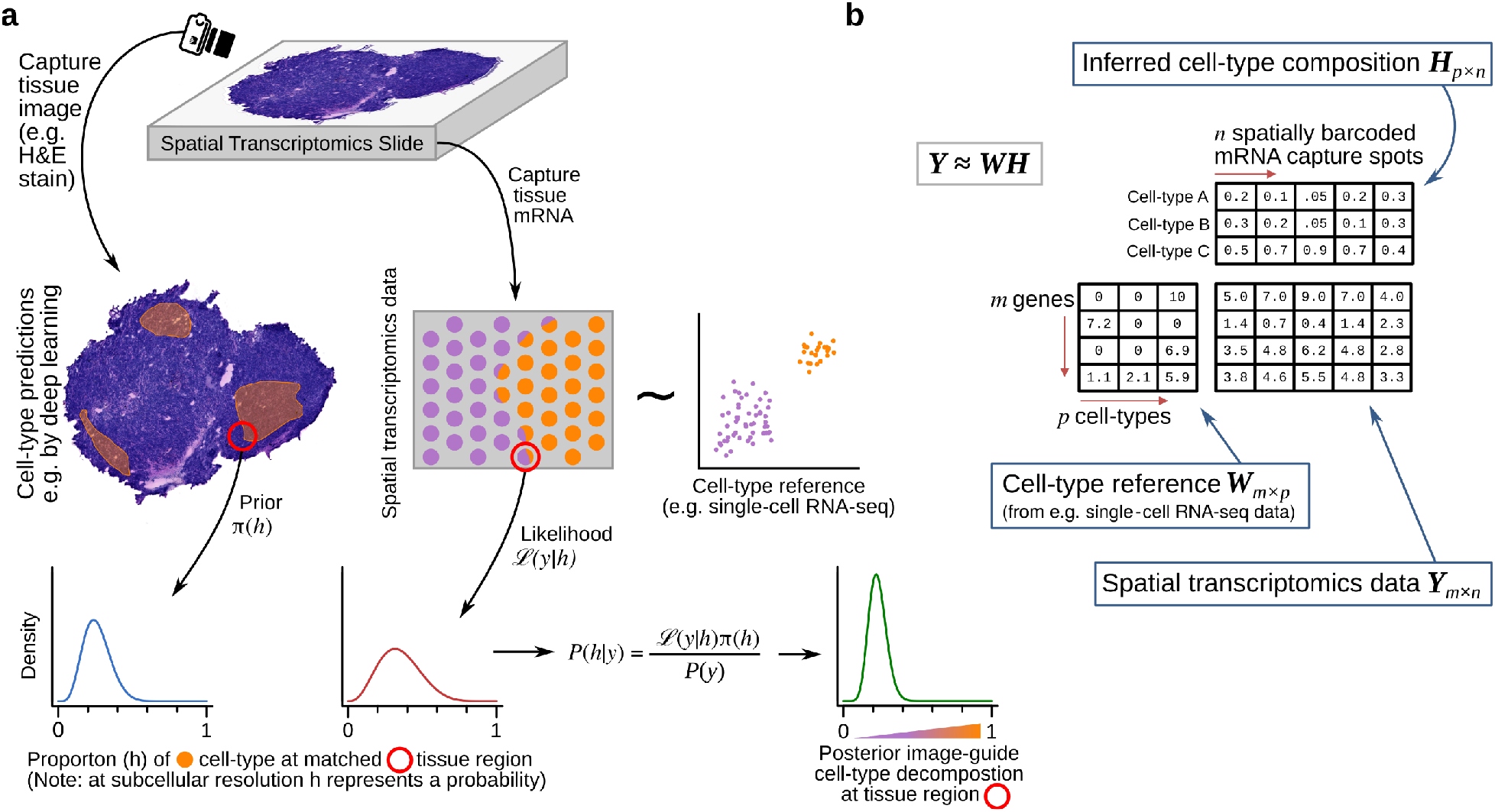
Schematic overview of Guiding-Image Spatial Transcriptomics (GIST) methodology. a) Schematic representation of GIST. The schematic shows a hypothetical tissue section, where we wish to identify the location of a hypothetical cell type (colored orange); this could represent, for example, immune cell infiltration in a tumor. Prior estimates of this cell type’s proportions from e.g. a deep learning model applied to an H&E stain image (left) are used to optimize the estimates derived from the spatial transcriptomics data (right), yielding improved estimates over what could be achieved otherwise (bottom right). b) Schematic representation of the cell type decomposition problem posed as a matrix decomposition. Spatial transcriptomics expression data is arranged in an *m* genes by *n* mRNA-capture-spots matrix ***Y***. This matrix is decomposed into a basis matrix ***W*** and a matrix ***H*** that contains the proportion of each of *p* cell types on each spot or (at subcellular resolution) the probability that a spot matches a cell type (shown for three hypothetical cell types A, B, and C). The basis matrix ***W*** is typically known and can be derived for example from single-cell RNA-seq data from the same or similar tissue. Given this, all existing cell type decomposition algorithms, be they designed specifically for spatial transcriptomics data or not, aim to estimate ***H***.

In the datasets used in this study, each element of ***W*** was derived from ψ, a reference single-cell RNA-seq dataset. Single-cell RNA-seq data is most often modelled using a negative binomial distribution (20) estimated for each gene *i*, in each cell type *k*, from the expression data of the available single-cells indexed by *l. Φ*_*i,k*_ represents the overdispersion parameter of such a distribution:

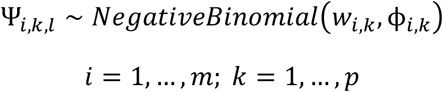

In this study, we approximated the elements of ***W*** by taking the mean normalized (details below) expression of each gene in each cell type in the reference single-cell RNA-seq dataset ψ, which in practical terms avoids having to include the entire single-cell RNA-seq dataset during the model fitting procedure, thus speeding up inference and likely having advantages on very large reference datasets.

Given ***Y*** and ***W***, the following model was then used for estimating ***H***:

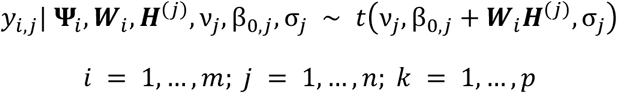

We placed a gamma prior (priors are denoted herein by π) on the degrees of freedom parameter *v* of the *t*-distribution, using shape and rate parameter values previously proposed by Juarez and Steele(21):

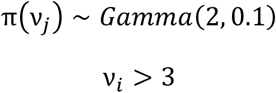

We constrained the elements of ***H*** to be positive and to sum to one within each spot:

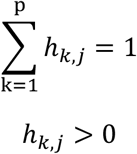

This was achieved by placing a non-informative Dirichlet prior on the columns of ***H***:

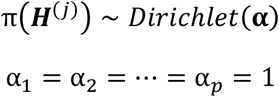

Each σ_*j*_ was assigned a non-informative prior.

We used the image data to generate a prior estimate of the abundance of some cell type *a* (e.g. immune cells) at each spot *j* (details below), then we placed a beta distribution prior on the corresponding proportion of cell type *N* at spot *j*:

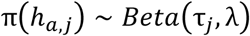

Here, τ_*j*_ is the mean of the beta distribution representing the image-derived prior estimate for the proportion of this cell type *a* at spot *j*. λ is a hyperparameter, representing the total count parameter of the beta distribution, determining how much weight is to be placed on the image data and how much to place on the transcriptomic data. Elements of ***H*** outside of the specific row-of-interest *a*, (i.e. elements for which an informative image-derived prior is not available) are assigned non-informative priors.

In the notation above, vectors are shown using boldface and matrices bold capital letters. We assume *m* genes (indexed by *i*), *n* spots, (indexed by *j*), and *p* cell types (indexed by *k*).

### Fitting the GIST and GIST base-model

The statistical model described above was implemented in the Stan programming language using the *rstan* package. The Hamiltonian Monte Carlo (HMC) algorithm was used to estimate the model parameters. The HMC algorithm was run for 2000 iterations where the first 1000 iterations were discarded as burn-in. The posterior mean was used as final parameter estimates.

### Prior construction

#### Mouse brain dataset

To avoid outlier bias in the IF image data the pixel-level image intensity values were first capped at the 99^th^ percentile and values below the 1^st^ percentile were set to zero. These pixel-level intensity values were then rescaled from 0 to 1, by dividing all values by the maximum capped value. Pixels overlapping each spatial transcriptomics mRNA capture spot were defined as those centered around the middle of the spatial transcriptomics spot in a 70-pixel radius—the center of the spot was defined in an annotation file that was output by the 10x Genomics SpaceRanger software. The rescaled pixel-level intensity values were then averaged over the slide regions corresponding to each spatial transcriptomics spot to obtain a single intensity value for each spot. This procedure was repeated for both IF channels—RBFOX3 (Neuron) and GFAP (Glia). Finally, the intensity values for each spot in each channel were mapped onto the quantiles of the cell type proportion estimates obtained from a first round of model fitting using the GIST base-model (i.e. where all parameters estimating cell-type abundance are assigned non-informative priors). These IF image-derived mapped spot level intensity values, which act as a proxy for the abundance of neurons or glia, were used as priors on the appropriate parameters in the GIST model.

#### Breast cancer dataset

The deep learning models used in the breast cancer analyses were previously published by Saltz *et al*. (22) and were obtained from the Quantitative Imaging in Pathology (QuIP) group’s website (https://sbu-bmi.github.io/quip_distro). These are convolutional neural network-based deep learning models, which had been pre-trained to recognize tumor-infiltrating lymphocytes. The original authors had trained these models using pathologist annotated H&E-stained tissues sections from TCGA. We used the VGG16-based model provided by the group. The breast cancer H&E images were converted from JPEG format to tiled TIFF format and the software suite VIPS was used to encode the TIFF files with a micron per pixel (MPP) value for each slide. The encoded TIFF files were processed using QuIP’s deep learning pipeline to generate a probability map over the entirety of each breast cancer H&E stained slide image. The deep learning model assigned probability values to patches of 50×50 microns. For a given spot, the assigned patch-level probability values were converted to spot-level probability values by taking a weighted sum of the patches, where the weight is the pixel overlap between the patch and the spot. This generated values for each spatial transcriptomics spot that approximately corresponded to the probability of immune cell infiltration. Similarly to the mouse brain IF dataset, these probability values were then mapped onto the distribution of total lymphocyte (T cell and B cell) content estimated from gene expression-derived proportions alone, obtained by an initial round of model fitting using the GIST base-model. These mapped values were used as informative priors on the appropriate model parameters in the GIST model.

The image processing code was implemented in Python using imaging libraries PIL.Image and imageio. Visualization and analysis of imaging data were carried out using the NumPy, pandas, and Matplotlib libraries.

### Quantifying the improvement achieved by the GIST model, compared to an expression-only model, by benchmarking against a pathologist-defined ground truth

For each slide in the breast cancer dataset, we quantified a model’s ability to accurately estimate regions of immune cells by the median of immune cell proportions in spots labeled as immune-infiltrated by the original pathologist, divided by the median of immune cell proportions estimated in the other remaining spots:

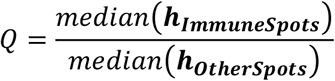

***h***_***ImmuneSpots***_ is a vector of model-estimated immune cell proportions for spots annotated by the pathologist as containing immune cells, and ***h***_***OtherSpots***_ are the immune cell proportions estimated at the other spots on the same slide.

With better performance, the scalar value *Q* will increase, as the model’s output better matches the pathologist-defined ground truth for this slide. Having defined this performance metric, we defined the improvement of the GIST model over the expression-only GIST base-model below as Δ, a scalar representing the difference between this ratio statistic *Q* when immune cell proportions were estimated with the GIST model (*Q*_*GIST*_) or the GIST base-model (*Q*_*GISTBa****se****M****o****d****e****l*_):

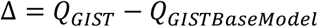

To assess whether the improved performance Δ observed for the GIST model over the GIST base-model was statistically significant, we used a permutation-based strategy, building a null distribution by randomly shuffling the pathologist’s spot level annotations. Specifically, for each permutation, the spots were randomly assigned as either immune infiltrated or non-immune, fixing the total number of immune infiltrated spots to the same number as the pathologist’s annotation of that slide; we then computed the improvement in the performance Δ_*perm*_ of the GIST model over the GIST base-model using the same procedure that was applied to the real arrangement of the pathologist’s annotations. This was repeated for 100,000 permutations, generating a null distribution against which to compare the observed test statistic Δ. A *P*-value was then calculated by the proportion of permuted values Δ_*perm*_ that achieved a value at least as extreme as Δ, the test statistic observed in the pathologist’s real annotations. In the cases where no permutated value more extreme than the original test statistic was observed (G1 and H1), a *P*-value was calculated by approximating the null distribution using a normal distribution, with a mean and standard deviation equal to that of the Δ_*perm*_ values from the 100,000 permutations. Note that while the statistical validation procedure described in this section assumes a pathologist’s manual annotation of the slides as the hold-out ground truth, these same procedures could be applied to any assumed ground truth annotation given any class of informative paired image, for example, substituting the pathologist’s annotation for an orthogonal IF or pathology stain marking the cell type of interest.

### Second pathologist’s re-annotation of the breast cancer spatial transcriptomics slides

A second pathologist was asked to assign new immune infiltration grades from H&E images of spots for three spatial transcriptomics breast cancer slides – B1, C1, and H1. The pathologist (co-author Dr. Heather Tillman) was asked to *blindly* score H&E images of slide regions overlapping the spatial transcriptomics mRNA capture spots from three groups of spots: These were (i) spots that were annotated as immune cell infiltrated by the original pathologist (slide H1 only), (ii) spots that were identified as high-confidence immune infiltrated by the GIST model, or (iii) other randomly chosen spots. High-confidence immune-cell-infiltrated spots from the GIST model were selected as the spots having a predicted proportion of immune cells that was greater than the upper quartile plus 1.5 times the interquartile range of the data, a *de facto* metric used to define outliers. For each slide, the number of random spots selected was equal to the number of spots included from the GIST model. This second pathologist was then asked to score/grade an H&E stain image of each spot, scoring immune cell infiltration levels as low, middle, or high, while remaining blinded to the group from which the spot image was selected. This provided a new score for each spot from each of the three groups (annotated, GIST, random). We then applied a one-sided Wilcoxon rank-sum test to assess whether these scores were significantly higher in the group of spots predicted as high confidence immune infiltrated by the GIST model compared to the randomly selected spots or the immune infiltrated spots from the initial pathologist’s annotation, where low, middle and high scores were encoded on an ordinal scale as 1, 2 and 3 respectively.

### Simulations to assess the ability of the GIST base-model to accurately identify cell type composition in gene expression data from a mixture of cell types

#### Splatter

The accuracy of cell type proportions estimated from the various computational methods was compared to the GIST base-model by first creating synthetic mixtures of gene expression data using the popular Splatter model (23). We used the Splatter model with a slight modification, which was recently proposed by Zhang *et al*. (24), who reported that the native Splatter model did not capture the empirical distribution of log fold changes observed in real data. The enhanced Splatter model was obtained from the GitHub repository of Zhang *et al*. (https://github.com/Irrationone/splatter), where the authors had learned the simulation parameters from the counts matrix of a publicly available PBMC single-cell RNA-seq dataset generated by 10X Genomics. The parameters for log fold changes were learned by fitting a truncated student’s t-distribution to the log fold changes between B cells and CD4 T cells in this same PBMC dataset.

Using the enhanced Splatter framework, we generated a dataset with 100 gene expression samples, each created from mixtures of cell types, along with a simulated paired reference single-cell RNA-seq dataset. The paired single-cell RNA-seq data were collapsed by their mean to create the required reference signature matrix ***W***, which was passed to each of the computational methods. Each expression mixture sample was generated by taking a weighted average of gene expression across 100 cells (generated independently of the reference single-cell RNA-seq data) from each of six synthetic cell types. Ground truth cell type proportions for the 100 simulated mixture samples were randomly generated from a flat Dirichlet distribution, where each cell type was assigned equal weight.

#### Immune cell deconvolution

We performed a second set of benchmarking simulations using the framework developed by Strum *et al*. (25), which rather than relying entirely on simulation, created a mixture gene expression dataset by computationally mixing real single-cell RNA-seq data, previously generated by Schelker *et al*. (26). In this benchmark, ground truth was established by mixing gene expression counts from 500 single-cells from each of eight immune cell types in known proportions and the simulated mixture was created by taking an average across cells. For the fairest comparison, we supplied each of the methods the LM22 cell type signature matrix(27) (corresponding to ***W*** in our notation herein), which is a signature matrix created by the developers of CIBERSORT that represents average gene expression values in each of 22 immune cell types. Note this was not possible for Stereoscope, which only accepts single-cell RNA-seq data as the reference input, from which it estimates the cell type signature matrix internally. Because the LM22 cell types do not have a strict one-to-one correspondence with the cell types annotated in Schelker *et al*., the results were mapped to the most relevant cell type using the same mappings previously employed by Strum *et al*.

In all simulations, the performance of each method was summarized by the mean absolute error (MAE), which is the average of the absolute value of the difference between each predicted cell type proportion and the known simulated ground truth proportion:

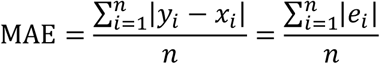

Where *y*_*i*_ is a predicted cell type proportion, *x*_*i*_ is the ground truth proportion, *e*_*i*_ is the error associated with the prediction, and *n* is the total number of predicted data points generated by a given method.

### Data preprocessing, filtering, normalization and imputation

All public datasets were obtained as preprocessed counts matrices, which had been processed according to the previous authors. Generally, spatial transcriptomics data displayed greater sparsity than the single-cell RNA-seq data, which arises because of differences in platform-specific mRNA capture efficiency. To alleviate this difference, we used a non-parametric imputation approach. Specifically, we used the knnSmooth (28) algorithm (available at the GitHub repository https://github.com/yanailab/knn-smoothing) to impute the spatial transcriptomics data.

For the IF mouse brain dataset, we set the “number of nearest neighbors to aggregate” parameter *k* to 5 and the “number of principal components” parameter *d* to 10 (author’s suggested default). For the breast cancer dataset, we used the same approach with slight modifications. The resolution of spots on the breast cancer slides was coarser than on the Visium array and transcript capture was poorer. Thus, to overcome these limitations, we combined the spots from all the breast cancer spatial transcriptomics slides and imputed them together using the knnSmooth algorithm with a *k* parameter of 10, mitigating the lower transcript capture efficiency in the breast cancer dataset.

Thereafter, both the spatial transcriptomics and single-cell RNA-seq data were normalized separately by using Seurat’s SCTransform (29), which importantly removes technical effects such as library size effects. We restricted the single-cell RNA-seq and spatial transcriptomics data to the intersection of their 2,000 most highly variable genes, yielding totals of 1,024 and 837 genes used for GIST model fitting in the mouse and breast cancer datasets respectively.

### Software and code availability

The GIST model has been made available as an R package, which can be obtained at: https://github.com/asifzubair/GIST

All the code for the analyses presented in this manuscript are available on GitHub: https://github.com/asifzubair/GIST-paper

## RESULTS

### Guiding-Image Spatial Transcriptomics (GIST) jointly leverages spatial transcriptomics and paired tissue images to improve cell type decomposition

GIST attempts to improve cell type decomposition in spatial transcriptomics data by leveraging prior estimates of cell type composition from paired tissue images. The method relies on Bayesian probabilistic modeling, a statistical approach that naturally lends itself to integrating multiple sources of information, jointly leveraging spatial transcriptomics and imaging information to improve cell type decomposition estimates. Intuitively, the approach uses the imaging data to provide an initial “suggestion” as to the cell types in a particular region of the spatial transcriptomics slide, but this suggestion can be overcome if outweighed by the evidence from the transcriptomic data (schematic representation in Fig. 1a, see Methods for technical details of model).

### A Bayesian probabilistic model for cell type decomposition performs competitively when compared to existing methods in simulations when no paired image information is leveraged

Existing methods for cell type decomposition in spatial transcriptomics data are related to previous models for bulk gene expression deconvolution and can be broadly conceptualized as a matrix decomposition, where some reference basis matrix of expression data from purified cells ***W*** (e.g. derived from single-cell RNA-seq) is used to estimate the proportion of each cell type ***H*** in the bulk mixture ***Y*** (Fig. 1b for schematic representation). At subcellular resolution, the ***H*** matrix can be thought of as probability estimates, rather than proportion estimates (15), although for simplicity we use the term “proportion” throughout this manuscript.

The statistical model underlying GIST is related to these existing approaches but includes the ability to leverage prior information derived from paired tissue images. Thus, we were first interested in assessing whether our model performed competitively when compared to existing approaches in the absence of prior information derived from images (*henceforth referred to as the “GIST base-model”*). To test this, we first developed two complementary unbiased benchmarking simulations, one based on the existing tool Splatter (23) and one based on a published benchmarking dataset (25), which evaluates methods on a simulated mixture of immune cell types from a real single-cell RNA-seq dataset. We compared the GIST base-model to two methods originally designed for bulk gene expression data (CIBERSORT (30), DeconRNASeq (31)), two methods tailored specifically for spatial transcriptomics data (Stereoscope (17) and SpatialDWLS (32)), and linear regression (the simplest conceivable model.) Based on the mean absolute error (*MAE*), CIBERSORT performed slightly better on the Splatter simulations (Fig. 2a, Supplementary Figure 1, Supplementary Table S1; *MAE* = 7.9 × 10^−2^ for CIBERSORT and 8.3 × 10^−2^ for the GIST base-model), while the GIST base-model performed best on the other benchmarking dataset (Fig. 2b, Supplementary Figure 2, Supplementary Table S2; *MAE* = 0.09 for CIBERSORT and 0.06 for the GIST base-model). However, given the conceptual similarity of the underlying models, it is not surprising that none of these existing methods produce markedly dissimilar results in either simulation, suggesting that, rather than further model tweaking and optimization, a new conceptual advance may be necessary to achieve meaningful progress on the cell type decomposition problem.

**Figure 2:**
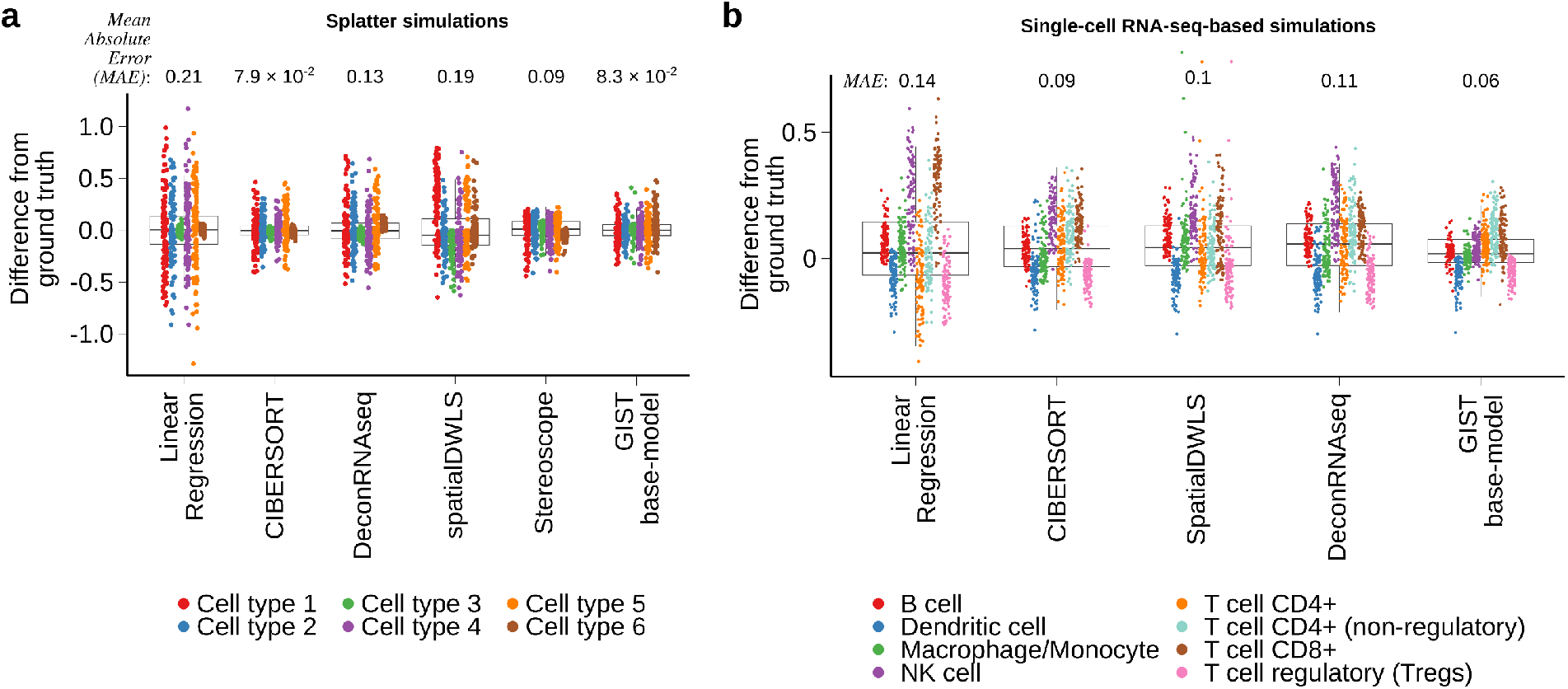
A Bayesian probabilistic model performs similarly to existing cell type decomposition methods when no prior information is available. a) Boxplot showing the results of five cell type decomposition methods on simulated mixture gene expression data, for a mixture of 6 cell types, generated using the tool Splatter (see Methods). Points have been colored by the simulated cell type and the y-axis shows the deviation from ground truth, quantified by the difference between the estimated cell type proportions in a sample and the true proportion used as ground truth for the simulation. The Mean Absolute Error (*MAE*), summarizing the overall performance of each method is as follows (lower values imply better performance): Linear regression = 0.21, CIBERSORT = 7.9 × 10^−2^, DeconRNAseq = 0.13, spatialDWLS = 0.19, Stereoscope = 0.9, GIST base-model = 8.3 × 10^−2^. b) Similar to (b) but based on the simulated dataset obtained from the benchmarking procedure outlined in Strum *et al*. (25). Points have been colored by the immune cell type and the y-axis shows the deviation from ground truth, quantified by the difference between the estimated cell type proportions in a sample and the true proportion used as ground truth for the simulation. The Mean Absolute Error (*MAE*), summarizing the overall performance of each method is as follows (lower values imply better performance): Linear regression = 0.14, CIBERSORT = 0.09, DeconRNAseq = 0.11, SpatialDWLS = 0.1, GIST base-model = 6.4 × 10^−2^. Note: Stereoscope was not included in this second set of simulations because it was not possible to pass the CIBERSORT LM22 signature matrix, which was used as the cell-type reference in this simulation, to Stereoscope (see Methods). In all boxplots, the center line represents the median, bound of box is upper and lower quartiles and the whiskers are 1.5× the interquartile range.

### The GIST base-model performs competitively on spatial transcriptomics data obtained from mouse brain sections when cell type specific immunofluorescence markers are treated as a ground truth

We were next interested in comparing the performance of the GIST base-model to other methods using real spatial transcriptomics data. To do this, we leveraged a publicly available dataset (see Data Availability), which measured gene expression in the mouse brain using the popular 10x Genomics Visium spatial transcriptomics platform, and where immunofluorescence (IF) staining was performed on the reverse side of the tissue section. These IF stains were conducted for two proteins, RBFOX3 and GFAP, which are protein markers unique to neurons and glia respectively (Fig. 3a). We calculated the average pixel intensity of each of these two markers in all image pixels overlapping each spatially barcoded mRNA capture spot on the Visium slide (Fig. 3b; see Methods), then we used these spot-level intensity estimates to represent an independent ground-truth approximating the abundance of neurons and glia in regions of the slide overlapping each of the Visium array’s 4,992 spots.

**Figure 3:**
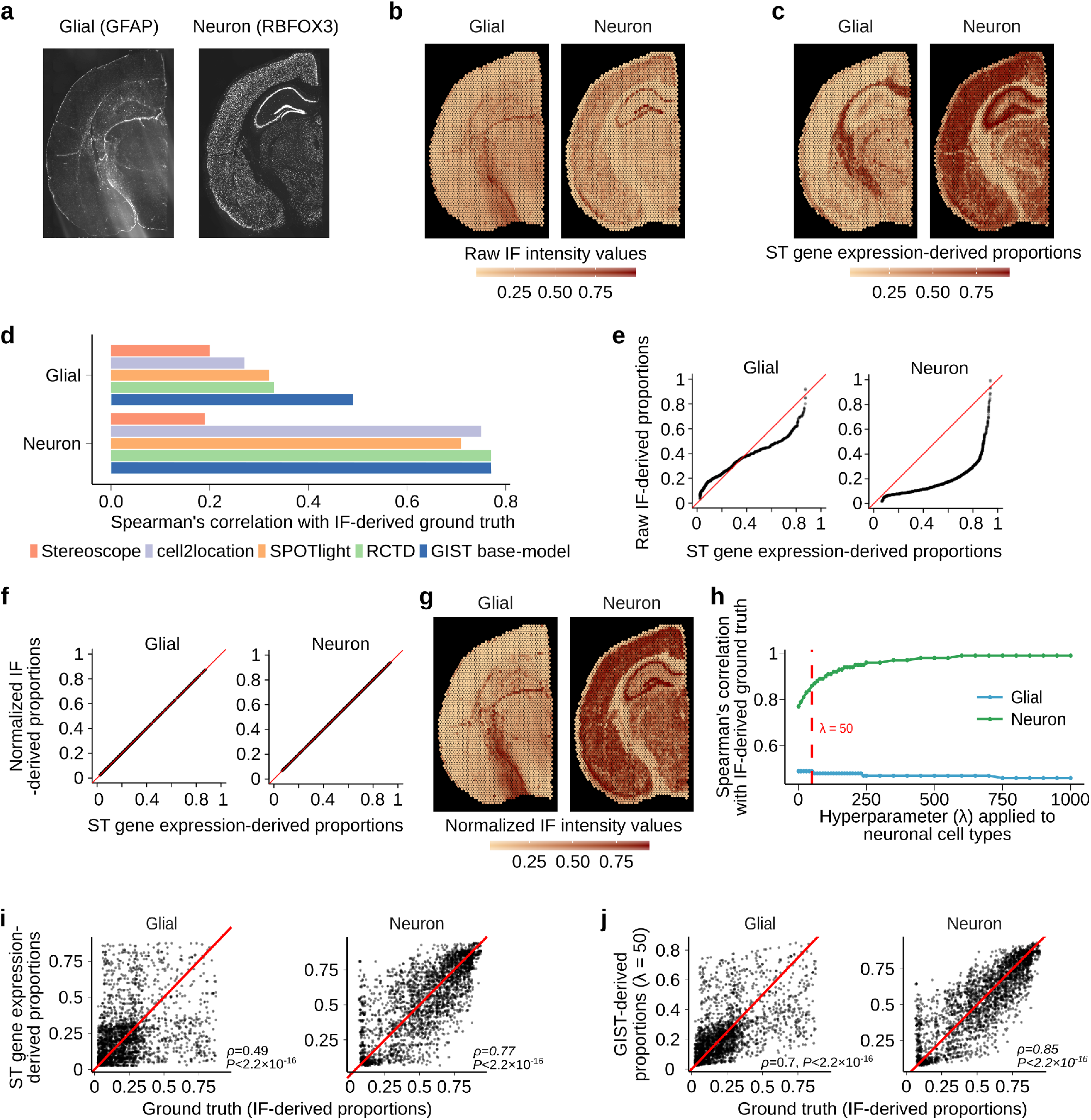
Incorporating image-derived prior information from matched immunofluorescence stains in mouse brain spatial transcriptomics data. a) Raw immunofluorescence image of the mouse brain tissue section showing the glial (GFAP) and neuronal (RBFOX3) cell markers. b) Spatial distribution of raw IF intensity values for GFAP (glial) and RBFOX3 (neuronal) when fluorescence intensity has been averaged over pixels corresponding to each spatial transcriptomics spot’s location. Intensity values were rescaled from 0 to 1. c) Spatial distribution of glial and neuronal proportions estimated from the spatial transcriptomics gene expression data using the GIST base-model. d) Bar plot showing Spearman’s correlation between IF-derived ground truth cell type proportions and cell type proportions estimated from five different gene expression-based spatial transcriptomics cell type decomposition methods (Stereoscope, cell2location, SPOTlight, RCTD, and the GIST base-model). e) Quantile-quantile plot (QQ plot) of image-based IF-derived values for total glial and neuronal content for each spot (y-axis) versus values obtained for total glial and neuronal content from the spatial transcriptomics gene expression data only using the GIST base-model (x-axis). f) Same as in (e) except that this QQ plot is generated after post-mapping normalization where the distribution of cell type compositional estimates from the IF images were mapped onto the distribution of cell type compositional estimates from the spatial transcriptomics gene expression data generated using the GIST base-model. g) Spatial distribution of IF intensity values for the glial and neuronal channel where the values have now been mapped to a distribution estimated from the gene expression data using the GIST base-model. h) Line plot showing the change in GIST model performance as we increase the key hyperparameter *λ* (x-axis). Performance is quantified by Spearman correlation with IF-derived ground truth (y-axis) and is shown for both neuronal (green) and glial (blue) cell types. The RBFOX3 IF image-derived prior is only applied to the neuronal cell type. A non-informative prior is applied to the glial cell type. The vertical dashed red line indicates a stopping point (*λ* = 50) where performance in the glial channel begins to deteriorate, indicating the model has been overfitted to the RBFOX3 IF data. i) Scatter plots showing the cell type compositional estimates against IF-derived ground truth (x-axis) in the mouse brain for glia (left) and neurons (right) derived from the spatial transcriptomics gene expression data using the GIST base-model (y-axis) when no prior information is leveraged. *P*-values are from Spearman’s correlation test. j) Similar to (i) but showing the improved agreement with ground truth (x-axis) when the IF-derived cell type compositional estimates are incorporated as prior information using the GIST model with a *λ* hyperparameter value of 50 (y-axis). *P*-values are from Spearman’s correlation test. Abbreviations: ST, Spatial Transcriptomics; IF: Immunofluorescence.

Next, using the GIST base-model, we estimated the cell type composition on each spot from the spatial transcriptomics data by leveraging a single-cell RNA-seq dataset that was available from a similar region of a mouse brain, allowing us to estimate the abundance of glial and neuronal cell types from the spatial transcriptomics expression data alone (Fig. 3c). We compared the results obtained from the GIST base-model to popular spatial transcriptomics cell type decomposition methods Spotlight (14), RCTD (15), Stereoscope (17), and cell2location (33), treating the IF-derived estimates of neurons and glia at each spot as ground truth. Consistent with our simulations, the GIST base-model, RCTD, cell2location, and Spotlight all performed quite similarly in these benchmarks on real data; however, we note that the GIST base-model had slightly better performance than the other methods, achieving Spearman’s rank correlations of 0.49 and 0.77, compared to 0.33 and 0.77 for RCTD (the second best performing method), for the glial and neuronal comparisons respectively (Fig. 3d; *P* < 2.2 × 10^−16^ from Spearman’s correlation against IF-derived ground truth for all five methods; Supplementary Figures 3-7). Overall, these results suggest that the GIST base-model performs competitively when compared to existing methods for cell type decomposition in real spatial transcriptomics data.

### Incorporating image-derived prior information from matched immunofluorescence stains has the potential to improve cell type decomposition in spatial transcriptomics data

Even though our GIST base-model performed well compared to existing methods, the results above also showed that the best-performing methods were not markedly different and fall well short of an optimal performance when compared to the IF-derived ground truth. Thus, we next hypothesized that it should be possible to markedly improve our performance by leveraging our model’s Bayesian implementation and supplying the model with informative image-derived prior information (*henceforth referred to as the “GIST model”*). We reasoned that we could first demonstrate this principle on this mouse brain dataset, leveraging the IF-derived estimates of cell type abundance. However, IF-derived pixel intensity estimates do not represent proportions on a 0-1 scale and thus it is not obvious how this information could be leveraged as prior estimates of cell type composition in the GIST model. To solve this problem, we first normalized the IF-derived estimates by mapping them onto the quantiles of the spatial transcriptomics-derived cell type proportion estimates, generated by an initial round of model fitting using the GIST base-model (Fig. 3e-g; see Methods). We then refit our GIST model, incorporating this prior knowledge derived from the RBFOX3 IF data, providing “suggestions” of the abundance of neuronal cell types over each spatial transcriptomics spot. We specified these priors using a beta distribution applied to the appropriate group of model parameters corresponding to neuronal cell type estimates. The beta distribution was parameterized by its mean (*τ*; the point estimate of the normalized cell type proportion estimate from the IF image) and the total-count parameter (*λ*; the strength of the prior, corresponding to the weight placed on the IF image)— any beta distribution is naturally constrained to a 0-1 scale, meaning it is appropriate for specifying image-derived prior estimates of cell type composition. The key modeling question is then determining how much weight to place on these image-derived priors and how much to place on the spatial transcriptomics data itself. This must be determined by selecting the hyperparameter *λ*, where a value that is too small will mean there is little to no influence of the image-derived cell type information on the model’s output, but selecting a value that is too large will cause the model to over rely on the image and degrade performance.

We chose this hyperparameter *λ* by observing how the estimates of glial cell type composition compared to IF-derived glial-cell ground-truth (GFAP stain) when fitting the model with ever-increasing values of *λ* for the IF-derived neuronal cell type prior (RBFOX3 stain), only placing priors on the neuronal cell types. As expected, when increasing the value of *λ* and placing more weight on the image-derived prior for neuronal cells, the model’s output progressively more closely matched these IF-derived estimates for the neuronal cell types (Fig. 3h). However, as we continued to increase *λ*, placing more and more weight on the image-derived estimates of neuronal cells, we eventually observed a monotonic drop-off in the model’s performance, as measured by the agreement between the glial cell type estimates from the GIST model and the IF-derived ground truth from the GFAP glial marker protein (Fig. 3h). This monotonic drop-off begins at *λ* ***=*** 50, suggesting that beyond this point this prior is too strong, providing us a stopping criterion and a reasonable initial value of *λ* for image-derived priors. This value of *λ* concentrates most of the prior probability mass within approximately ±10% of the mean. At this *λ* value, the Spearman’s rank correlation between the model-derived neuronal cell type estimates and the IF-derived ground truth increased from 0.7 to 0.85, but this increase would be expected given that the model’s posterior predictions are being compared to the specified prior (Figs. 3i and 3j). While these analyses specify a statistical model (including reasonable hyperparameter estimates) and normalization procedure for directly leveraging cell type informative image-derived information, determining whether this improves performance would involve comparison against an independent data modality, which is not available for this dataset. We address this problem below, leveraging this initial estimate of the key hyperparameter *λ* in a new out-of-batch independent dataset and assessing the model performance against a pathologist manual annotation of paired H&E stained pathology images, which are available in this additional dataset.

### Incorporating prior information derived from deep learning models applied to matched H&E-stained images improves estimates of immune cell infiltration in breast cancer spatial transcriptomics data

The results above suggest it should be possible to improve cell type decomposition in spatial transcriptomics data by leveraging matched images. However, while IF stains can provide reliable markers of cell types, they are restricted to a small number of proteins and are currently less commonly collected than the H&E stain. Thus, we also wondered whether it would be possible to leverage cell-type-informative information derived from deep learning models applied to H&E stains—the principal pathology stain that is collected as a part of almost all sequencing-based spatial transcriptomics protocols. Deep learning models have already been developed that can output numerous clinically relevant annotations from H&E-stained tissue section images alone, which could theoretically be usefully propagated in the spatial transcriptomics assay. These annotations include cell type composition, expression of signaling pathways, chromosomal ploidy, and immune cell infiltration (22,34,35). To test whether such information could be usefully exploited in spatial transcriptomics assays, we obtained 8 previously published spatial transcriptomics tissue slides, which had measured gene expression in biologically independent breast cancer tumors. Critically, each of these tissue sections had also been H&E stained (Fig. 4a, panel (a) in Supplementary Figures S8-S12), and regions of immune cell infiltration had been annotated by a previous pathologist (Fig. 4b, panel (b) in Supplementary Figure S8-S12), providing an independent ground truth against which to assess our model predictions (notably this was not available for the previous IF dataset). Identifying immune cell infiltration has prognostic value (36) and is predictive of response to cancer immunotherapy (37), hence represents a particularly interesting use case of the GIST model.

**Figure 4:**
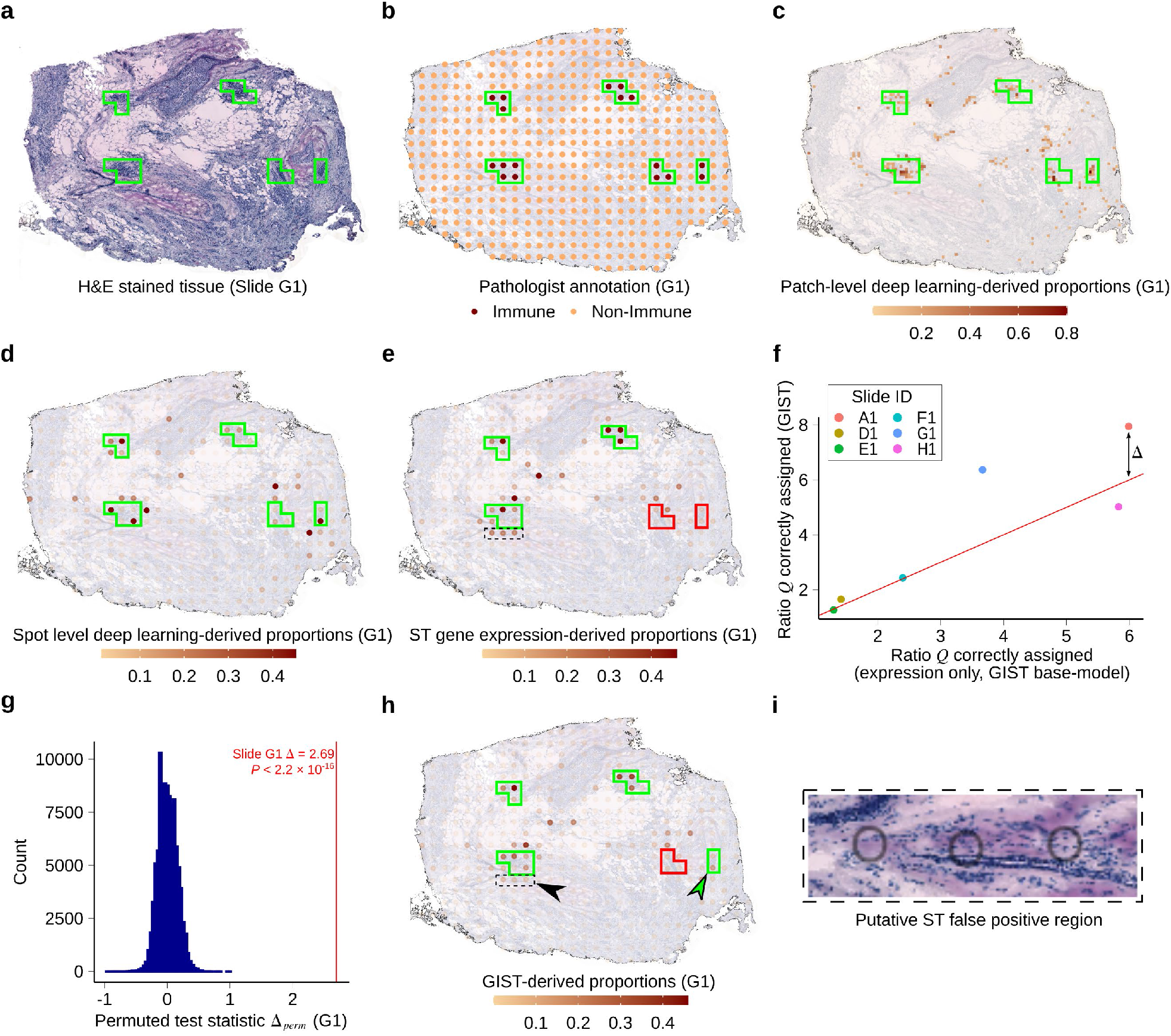
Tissue image-derived cell type compositional estimates can be leveraged to improve estimates of immune cell infiltration in breast cancer tissue sections profiled using spatial transcriptomics. a) H&E stained tissue image obtained from the reverse side of the breast cancer spatial transcriptomics slide G1. Green outline shows regions containing ST spots annotated as containing immune cells by the pathologist. b) Pathologist annotation for slide G1 showing regions containing spatial transcriptomics spots that were labeled immune cell infiltrated (marked by dark-colored spots and green outlines). c) Output from the deep learning model for slide G1 overlayed on top of the breast cancer tissue section H&E image. The color scale indicates deep learning-derived predictions for the proportions of immune cells made on 50×50 micron patches of the tissue. Green boxes outline regions of pathologist’s annotated immune spots. d) Slide G1 showing the patch level deep learning predictions converted to spot level predictions, so that they can be used as priors in the GIST model. Spot level predictions are a sum of patch level predictions weighted by their percent overlap with the spot. Boxes outline regions of pathologist’s annotated immune spots. e) Slide G1 showing the gene expression-derived immune cell proportions from the GIST base model. Solid boxes indicate the regions of the pathologist’s annotated immune spots. Green indicates that the model reasonably identifies immune-infiltrated spots. Red indicates that the immune spots were not captured by the model. The dashed black box indicates a region of interest that likely is a false positive (see panels (h) and (i)). f) Scatterplot showing the performance of the GIST model (y-axis) versus the performance of a base-model based on only gene expression data (x-axis) for six pathologist-annotated spatial transcriptomics slides. Performance is defined as the ratio of the median proportion of immune cells in pathologist labeled immune cell slide spots, versus the median proportion of immune cells in the other slide spots (*Q*, see Methods). Points are colored by slide ID. The red line is the identity line (intercept of 0, slope of 1), and the distance between this line and each point (black arrow) represents the observed test statistic Δ for that sample. g) Histogram showing the empirical null distribution of ratio-based test statistic (Δ_*perm*_, see Methods) generated using a permutation procedure (x-axis). The test statistic is a measure of improvement in model performance, versus the pathologist-annotated ground truth, when deep-learning derived prior cell type annotations are incorporated. The observed test statistic Δ is shown using a vertical red line. *P*-value from permutation test. h) Slide G1 showing the GIST model-derived immune cell proportions, when the deep learning immune cell type annotation has been used as an informative prior. Solid boxes indicate regions of pathologist’s annotated immune spots. Green indicates that immune spots were successfully identified, and red indicates that immune spots were not well captured. The dashed black box, highlighted by the black arrowhead, indicates the same region of interest as in (e), where the false positive immune cell predictions have been mitigated. The green arrowhead highlights a region where the correct identification of a pathologist annotated immune-infiltrated region has improved. i) Tissue image showing the region of interest highlighted by a dashed black box in panels (e) and (h). The H&E stain shows minimal evidence of immune infiltration in the areas overlapping the three spatial transcriptomics spots, whose location is shown by black circles. Abbreviations: ST, Spatial Transcriptomics.

Thus, we applied a previously published deep convolutional neural network (22), which had been trained using images collected as part of TCGA to identify regions of tumor-infiltrating lymphocytes from H&E stained tumor tissue sections. This yielded patches of deep learning-derived predictions of immune cell infiltration across each of our breast cancer tumor tissue sections (Fig. 4c, panel (c) in Supplementary Figures S8-S12), where gene expression had also been measured using spatial transcriptomics. We then averaged these deep learning derived predictions over the pixels overlapping each of the spatial transcriptomics mRNA capture spots, yielding a deep-learning-derived per-spot estimate of immune cell composition in each tumor (Fig. 4d, panel (d) in Supplementary Figures S8-S12, similar to the approach applied above for IF data; see Methods). Initial immune cell proportions at each spot were then estimated using the GIST base-model (Fig. 4e, panel (e) in Supplementary Figures S8-S12). We applied a similar normalization approach as we described for the IF data, mapping the deep learning derived estimates to the quantiles of the initial gene expression derived estimates, then applied these deep-learning-derived immune cell compositional estimates as informative priors, again specified as a beta distribution on the appropriate GIST model parameters. We used a *λ* value of 50, which was derived from the previous independent dataset (Fig. 3h). If the GIST model performs better than the expression-only GIST base-model, the expectation is that we should identify more immune cells in pathologist-annotated immune cell regions, but less in other regions of the slides. Thus, we quantified model performance by the ratio of immune cells identified within the pathologist’s annotated regions of immune infiltration, compared to all other regions of the tissue slide (this ratio is defined herein as *Q* (see Methods); note that regions of immune cells had been identified by the pathologist in six of eight slides). When compared to the pathologist-derived ground truth, the GIST model, leveraging deep learning-derived prior information, performed better than the expression-only GIST base-model in four out of the six slides (Fig. 4f, panel (f) in Supplementary Figures S8-S12). The performance increase over the GIST base-model was particularly large for two slides (Fig. 4g, panel (g) in Supplementary Figures S8-S12; increase in *Q* for GIST vs GIST base-model (defined herein as Δ) of 1.95 and 2.69, *P* = 7.2 × 10^−3^ and *P* < 2.2 × 10^−16^ for slides A1 and G1 respectively; empirical *P*-values were calculated by permutation, see Methods). Visual inspection of the results revealed examples of clear regions where leveraging the deep learning-derived prior information correctly decreased the estimates of immune cell composition in regions where the pathologist marked an absence of immune cells (Fig. 4h, black arrowhead, and Fig. 4i) and regions where estimates of immune cell composition increased to match the pathologist (Fig. 4h, green arrowhead). Given that the false positive region highlighted in Fig. 4i is directly adjacent to a true positive region of immune cells, it is plausible this represents diffusion of immune cell mRNA from the adjacent spots, a common issue in spatial transcriptomics assays that can seemingly be mitigated by jointly leveraging the paired tissue image. Thus, leveraging deep learning derived prior information has the potential to markedly improve cell type decomposition in data generated from spatial transcriptomics technologies.

### The GIST model identified large regions of immune cell infiltration that were missed by the initial pathologist

Surprisingly, one of the six breast cancer slides assessed demonstrated a statistically significant *decrease* in performance when we leveraged the image-derived prior estimates of immune cell infiltration (slide H1 in Fig. 4f, *P* = 3.56 × 10^−11^, Supplementary Figure S12g). However, closer inspection of this slide’s results revealed that there was a large region of this tumor that was identified as immune cell infiltrated by both the spatial transcriptomics assay and the deep learning model, but this region was not marked by the initial pathologist’s annotation (Supplementary Figure S12a-S12e and S12h)). Unsurprisingly, this region was predicted as heavily immune cell infiltrated by the GIST model, which also correctly identified the original pathologist’s annotated regions of immune infiltration in this slide (Fig. 5a, Supplementary Figure S12f).

**Figure 5:**
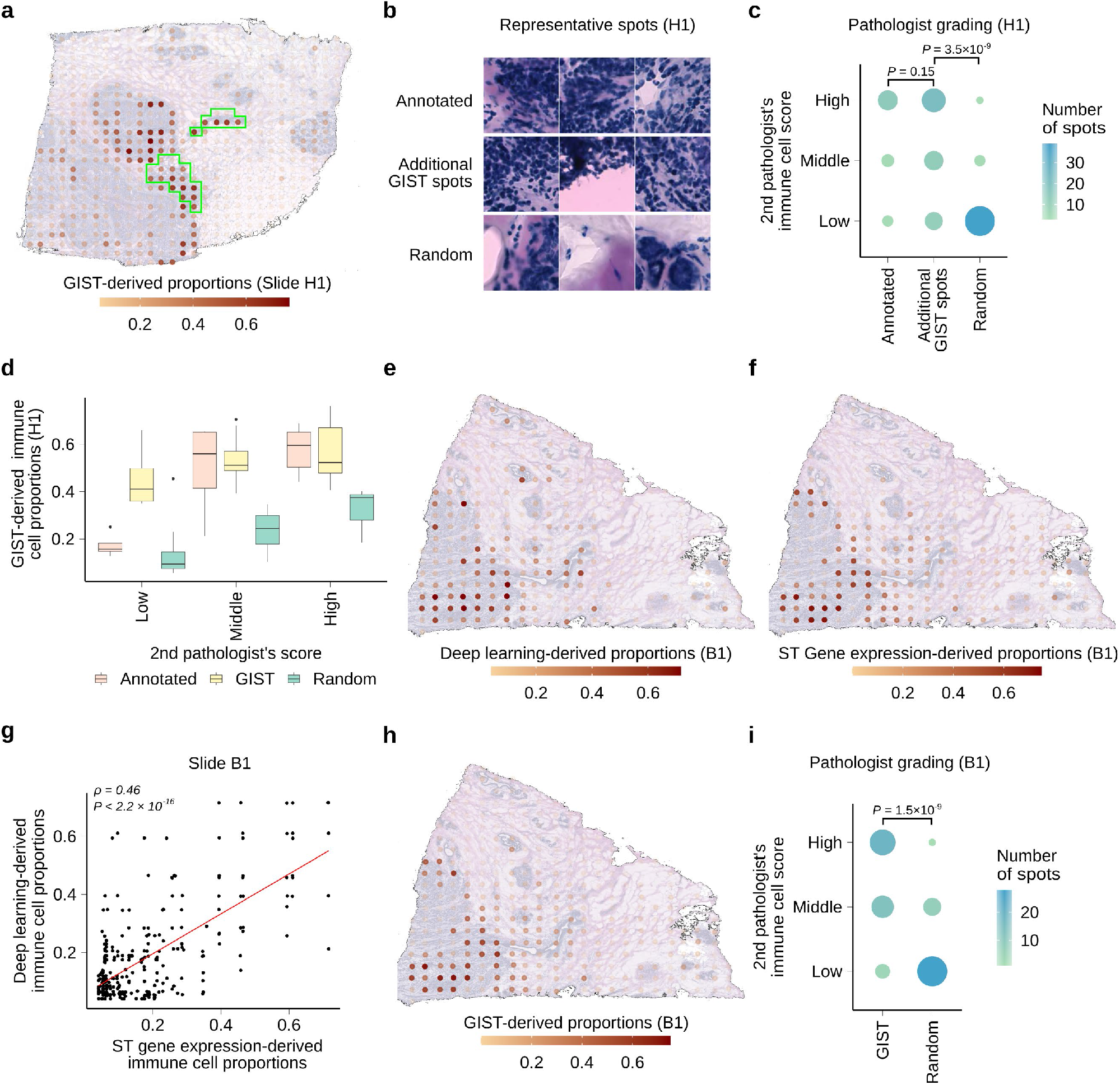
GIST model identifies regions of immune cell infiltration that were missed by an initial pathologist’s annotation. a) GIST model-derived proportions plotted on top of tissue from slide H1. Green outline indicates the original annotation of immune infiltrated spot regions identified by the initial pathologist. b) Three representative 100×100 micron images showing spots from the first pathologist’s annotated regions of immune cell infiltration (top), additional high confidence immune infiltrated regions identified by the GIST model (middle), and additional randomly selected regions (bottom). Spots are taken from slide H1. c) Dot plot showing the second pathologist’s immune infiltration grading with a score of low, middle, and high (y-axis) for spots from different regions of the tissue (x-axis). Spots were taken from slide H1 from regions previously annotated by the first pathologist as immune-rich, additional high confidence regions from the GIST model, and additional random regions on the slide. *P*-values from one-sided Wilcoxon rank sum test. d) Boxplot showing distribution of GIST model predicted immune cell proportions (y-axis) broken down by immune infiltration grade (x-axis) provided by the second pathologist. For each pathologist grade (low, middle & high), GIST scores are shown for spots from annotated, GIST high confidence, and random regions. Spots taken from slide H1. e) Deep learning-derived proportions for spots on slide B1. The color scale shows the predicted proportion of immune cells at a spot. f) Gene expression-derived proportions for slide B1 from GIST base-model. The color scale shows the predicted proportion of immune cells at a spot. g) Scatter plot showing the per-spot correlation between deep learning-derived predictions (y-axis) and ST gene expression-derived proportions (x-axis) for slide B1. Each dot is a spot and the red line is the regression line. *P*-value from Spearman’s correlation test. h) GIST model-derived proportions for slide B1. The color scale shows the predicted proportion of immune cells at a spot. i) Dot plot showing the second pathologist’s immune infiltration grading with a score of low, middle, and high (y-axis) for spots from different regions of the tissue (x-axis). Spots were taken from slide B1 from high confidence regions from the GIST model and random regions on the slide. *P*-value from one-sided Wilcoxon rank sum test. In all boxplots, the center line represents the median, bound of box is upper and lower quartiles and the whiskers are 1.5× the interquartile range. Abbreviations: ST, Spatial Transcriptomics.

Thus, we hypothesized that the apparent decrease in performance may have represented an oversight in the initial pathologist’s annotation, and thus a deficiency in the assumed ground truth, rather than a deficiency in the GIST model’s prediction. To test this, we devised a procedure that would allow a second independent pathologist (see Author’s Contributions) to re-examine the relevant regions of this slide, while remaining blinded to the GIST model’s output and the original pathologist’s annotation. The second pathologist was presented with (*n* = 115) 100 × 100-micron subregions from this slide and asked to categorize them as either low, middle, or high levels of immune cell infiltration. These subregions were chosen either from (i) the first pathologist’s annotated immune cell regions (ii) high-confidence immune cell regions identified by the GIST model but not the first pathologist or (iii) other randomly chosen regions (representative examples shown in Fig. 5b; see Methods). Remarkably, the second pathologist’s reannotation determined no statistical difference between the high-confidence regions of immune cell infiltration annotated by the first pathologist and the additional high-confidence regions identified by the GIST model, which were missed by the first pathologist (Fig. 5c; *P* = 0.15 from two-sided Wilcoxon rank-sum test). However, the high-confidence regions of immune cell infiltration identified by GIST were much more likely to be marked as high probability regions of immune cell infiltration when compared to randomly chosen slide regions (Fig. 5c, *P* = 3.5 × 10^−9^ from one-sided Wilcoxon rank-sum test). Additionally, the second pathologist’s high confidence immune infiltrated regions were mirrored by higher estimated proportions by GIST (Fig. 5d). These results support the notion that the additional regions identified by the GIST model were true regions of immune cell infiltration and that the poor performance on this slide arose from an omission in the original pathologist’s annotation, not falsely identified regions by the GIST model.

We also reexamined the two available spatial transcriptomics slides where the original pathologist’s annotation of the H&E images had not identified any regions of immune cell infiltration (Supplementary Figure S13a-b). Surprisingly, for both slides the deep learning model (Fig. 5e, Supplementary Figure 13c) and the expression-only cell type predictions from the spatial transcriptomics assay (Fig. 5f, Supplementary Figure 13d) agreed that there were in fact regions of immune cell infiltration (Fig. 5g, Spearman’s correlation = 0.46, *P* < 2.2 × 10^−16^; Supplementary Figure 13e, Spearman’s correlation = 0.25, *P* < 2.2 × 10^−4^). Unsurprisingly, these same regions were identified by the GIST model (Fig. 5h, Supplementary Figure 13f) and thus it seemed plausible that the initial pathologist had also missed these immune infiltrated regions in their initial examination of these two slides. We used the same scoring procedure outlined above to reannotate these slides by the second pathologist, who convincingly annotated these predicted regions as true regions of immune cell infiltration (Figure 5i, *P* = 1.5 × 10^−9^; Supplementary Figure 13g, *P* = 4.5 × 10^−2^; see Methods), which were also mirrored by higher proportions estimated by GIST (Supplementary Figure 13h-i).

Taken together, the presented use cases suggest that our GIST model, which can jointly leverage image-derived cell type annotations with spatial transcriptomics data, has the potential to improve cell type decomposition in spatial transcriptomics data and that such a strategy can be used to identify predictive and prognostically important features in human tissue sections.

## DISCUSSION

We have presented a conceptually novel computational methodology that can leverage cell-type-informative data derived from paired tissue images to improve inferences of cell type composition in spatial transcriptomics data. For the spatial transcriptomics platforms used in this study, these images were obtained from the reverse side of the slide-affixed tissue section (schematic in Fig. 1(a)), but it is also likely feasible to obtain informative images from an adjacent tissue section. One exciting application of the methodology is the ability to leverage inferences from deep-learning models applied to tissue images, for example H&E stains, which itself has recently reached close to pathologist level performance in annotating clinically relevant features of tissue sections (22,34,35). However, the statistical methodology is highly generalizable and could be applied to any image-derived prior information, which we have demonstrated for immunofluorescence. We have also described a set of generalizable statistical procedures for determining the value-added by incorporating paired cell-type-informative tissue images in spatial transcriptomics analysis (see Methods). These procedures were applied to our detailed example of leveraging deep learning-derived immune cell predictions in breast cancer, where a pathologist’s annotation was held-out as an assumed ground truth, but this same validation procedure could be applied to any imaging modality where a reasonable hold-out annotation is available, for example, in the context of IF data, derived from an orthogonal IF stain marking the cell type of interest. Generally, our proposed integrated approach has the potential to improve all downstream applications of spatial transcriptomics that rely on accurate cell type annotations, including identification of cell-cell or gene-gene interactions, or cell type specific differential expression (19).

Our framework will also spur the development of future similar computational approaches. Theoretically, any cell type decomposition method that could be re-implemented in a Bayesian framework could be adapted to leverage image-derived cell-type-informative prior information and this is likely possible for most of the existing models used in our comparisons-of-methods (Figs. 2 and 3). Thus, there is enormous scope for future model development and optimization within our novel framework. We also anticipate that our framework will lead to new modes of spatial transcriptomics experimental design. For example, we showed that immunofluorescence data could be leveraged. This opens the possibility of *a priori* staining for a few particularly informative protein markers, knowing that such markers can be used in downstream analyses to directly influence and improve the results of the spatial transcriptomics data analysis. This may be particularly useful for separating cell types when multicollinearity affects the performance of conventional models for cell type decomposition (18).

Additionally, while we have shown some illustrative examples, the Bayesian implementation allows enormous flexibility in how prior information is specified. It is theoretically possible to, for example, apply one prior to groups of cell types, or apply multiple partially overlapping priors derived from various sources of information. For the breast cancer dataset shown, we also fixed the *λ* hyperparameter to 50, using information obtained in the previous dataset. This is likely a conservative means by which to choose this key value and also assumes that this hyperparameter should be assigned the same value at all regions of the slide—almost certainly an oversimplification. Methods could likely be devised to adaptively adjust the value of the *λ* hyperparameter on a per-spot or per-slide-region basis, such that, for example, the differences in uncertainty associated with the deep learning-based outputs could be accounted for at each tissue region. Additionally, given that spatial transcriptomics is still a very new technology, the availability of datasets upon which to properly benchmark or test the GIST approach is still quite limited. Thus, particularly with increasing data availability, it is likely that creative applications within the described framework will eventually yield improvements over the results presented here.

In conclusion, we anticipate that jointly leveraging spatial transcriptomics and cell-type-informative images collected from the same or adjacent tissue sections will represent an important conceptually novel computational methodology, which has the potential to improve many applications of emerging spatial transcriptomics technologies, including potential translational applications in clinical and diagnostic pathology.

## DATA AVAILABILITY

### Mouse Brain

The mouse brain spatial transcriptomics Visium data with associated IF images were downloaded from the 10X Genomics website: https://support.10xgenomics.com/spatial-gene-expression/datasets/1.1.0/V1_Adult_Mouse_Brain_Coronal_Section_2

As a cell type reference ***W*** for these data, we used the curated mouse brain single-cell RNA-seq data provided by Andersson *et al*.^(17)^. This data had been originally retrieved from http://www.mousebrain.org and was processed by Andersson *et al*. for use in spatial transcriptomics analysis: https://github.com/almaan/stereoscope/tree/master/data/mousebrain

### Breast Cancer

The eight separate breast cancer spatial transcriptomics slides, previously generated by Andersson *et al*., were downloaded from https://github.com/almaan/her2st. This repository contained count matrices generated from the spatial transcriptomics assays, H&E images of the tissue sections (with and without pathologist annotation), and matrices detailing the location of the spots.

The single-cell RNA-seq breast cancer dataset, used to generate the cell type reference matrix ***W*** for all breast cancer analyses, was previously generated by Karaayvaz *et al*.(38) and obtained from: https://github.com/Michorlab/tnbc_scrnaseq.

## AUTHOR CONTRIBUTIONS

P.G. conceived and directed the project. A.Z. and P.G. wrote the code, with additional input from R.C. P.G. and A.Z wrote the manuscript. H.T. blindly re-annotated the breast cancer pathology slides to resolve the discrepancies between the GIST model and the original pathologist’s annotations. S.N., W.C.W, M.P., H.M.L, and J.E. provided additional support in data analysis and interpretation. All authors edited and approved the final manuscript.

## ACKNOWLEDGEMENTS

PG is supported by an NIGMS R35 award (R35GM138293) an R01 grant from NCI (R01CA260060), and a K99/R00 (R00HG009679) from NHGRI. PG also receives support from ALSAC.

## REFERENCES

1. Vickovic, S., Eraslan, G., Salmén, F., Klughammer, J., Stenbeck, L., Schapiro, D., Äijö, T., Bonneau, R., Bergenstråhle, L. and Navarro, J.F. (2019) High-definition spatial transcriptomics for in situ tissue profiling. Nature methods, 16, 987–990.

2. Stickels, R.R., Murray, E., Kumar, P., Li, J., Marshall, J.L., Di Bella, D.J., Arlotta, P., Macosko, E.Z. and Chen, F. (2021) Highly sensitive spatial transcriptomics at near-cellular resolution with Slide-seqV2. Nature biotechnology, 39, 313–319.

3. Rodriques, S.G., Stickels, R.R., Goeva, A., Martin, C.A., Murray, E., Vanderburg, C.R., Welch, J., Chen, L.M., Chen, F. and Macosko, E.Z. (2019) Slide-seq: A scalable technology for measuring genome-wide expression at high spatial resolution. Science, 363, 1463–1467.

4. Liu, Y., Yang, M., Deng, Y., Su, G., Enninful, A., Guo, C.C., Tebaldi, T., Zhang, D., Kim, D. and Bai, Z. (2020) High-spatial-resolution multi-omics sequencing via deterministic barcoding in tissue. Cell, 183, 1665–1681. e1618.

5. Chen, A., Liao, S., Ma, K., Wu, L., Lai, Y., Yang, J., Li, W., Xu, J., Hao, S. and Chen, X. (2021) Large field of view-spatially resolved transcriptomics at nanoscale resolution. bioRxiv.

6. Van de Velde, L.-A., Allen, E.K., Crawford, J.C., Wilson, T.L., Guy, C.S., Russier, M., Zeitler, L., Bahrami, A., Finkelstein, D. and Pelletier, S. (2021) Neuroblastoma formation requires unconventional CD4 T cells and myeloid amino acid metabolism. bioRxiv.

7. Moncada, R., Barkley, D., Wagner, F., Chiodin, M., Devlin, J.C., Baron, M., Hajdu, C.H., Simeone, D.M. and Yanai, I. (2020) Integrating microarray-based spatial transcriptomics and single-cell RNA-seq reveals tissue architecture in pancreatic ductal adenocarcinomas. Nature biotechnology, 38, 333–342.

8. Svensson, V., Teichmann, S.A. and Stegle, O. (2018) SpatialDE: identification of spatially variable genes. Nature methods, 15, 343–346.

9. Sun, S., Zhu, J. and Zhou, X. (2020) Statistical analysis of spatial expression patterns for spatially resolved transcriptomic studies. Nature methods, 17, 193–200.

10. Pham, D.T., Tan, X., Xu, J., Grice, L.F., Lam, P.Y., Raghubar, A., Vukovic, J., Ruitenberg, M.J. and Nguyen, Q.H. (2020) stLearn: integrating spatial location, tissue morphology and gene expression to find cell types, cell-cell interactions and spatial trajectories within undissociated tissues. bioRxiv.

11. Maaskola, J., Bergenstråhle, L., Jurek, A., Navarro, J.F., Lagergren, J. and Lundeberg, J. (2018) Charting tissue expression anatomy by spatial transcriptome decomposition. BioRxiv, 362624.

12. Tanevski, J., Gabor, A., Flores, R.O.R., Schapiro, D. and Saez-Rodriguez, J. (2020) Explainable multi-view framework for dissecting inter-cellular signaling from highly multiplexed spatial data. BioRxiv.

13. Arnol, D., Schapiro, D., Bodenmiller, B., Saez-Rodriguez, J. and Stegle, O. (2019) Modeling cell-cell interactions from spatial molecular data with spatial variance component analysis. Cell reports, 29, 202–211. e206.

14. Elosua-Bayes, M., Nieto, P., Mereu, E., Gut, I. and Heyn, H. (2021) SPOTlight: seeded NMF regression to deconvolute spatial transcriptomics spots with single-cell transcriptomes. Nucleic acids research, 49, e50–e50.

15. Cable, D.M., Murray, E., Zou, L.S., Goeva, A., Macosko, E.Z., Chen, F. and Irizarry, R.A. (2021) Robust decomposition of cell type mixtures in spatial transcriptomics. Nature Biotechnology, 1-10.

16. Biancalani, T., Scalia, G., Buffoni, L., Avasthi, R., Lu, Z., Sanger, A., Tokcan, N., Vanderburg, C.R., Segerstolpe, A. and Zhang, M. (2020) Deep learning and alignment of spatially-resolved whole transcriptomes of single cells in the mouse brain with Tangram. bioRxiv.

17. Andersson, A., Bergenstråhle, J., Asp, M., Bergenstråhle, L., Jurek, A., Navarro, J.F. and Lundeberg, J. (2020) Single-cell and spatial transcriptomics enables probabilistic inference of cell type topography. Communications biology, 3, 1–8.

18. Li, B., Liu, J.S. and Liu, X.S. (2017) Revisit linear regression-based deconvolution methods for tumor gene expression data. Genome biology, 18, 1–5.

19. Cable, D.M., Murray, E., Shanmugam, V., Zhang, S., Diao, M.Z., Chen, H., Macosko, E., Irizarry, R.A. and Chen, F. (2021) Cell type-specific differential expression for spatial transcriptomics. bioRxiv.

20. Sarkar, A. and Stephens, M. (2021) Separating measurement and expression models clarifies confusion in single-cell RNA sequencing analysis. Nature Genetics, 53, 770–777.

21. Juárez, M.A. and Steel, M.F. (2010) Model-based clustering of non-Gaussian panel data based on skew-t distributions. Journal of Business & Economic Statistics, 28, 52–66.

22. Saltz, J., Gupta, R., Hou, L., Kurc, T., Singh, P., Nguyen, V., Samaras, D., Shroyer, K.R., Zhao, T. and Batiste, R. (2018) Spatial organization and molecular correlation of tumor-infiltrating lymphocytes using deep learning on pathology images. Cell reports, 23, 181–193. e187.

23. Zappia, L., Phipson, B. and Oshlack, A. (2017) Splatter: simulation of single-cell RNA sequencing data. Genome biology, 18, 1–15.

24. Zhang, A.W., O’Flanagan, C., Chavez, E.A., Lim, J.L., Ceglia, N., McPherson, A., Wiens, M., Walters, P., Chan, T. and Hewitson, B. (2019) Probabilistic cell-type assignment of single-cell RNA-seq for tumor microenvironment profiling. Nature methods, 16, 1007–1015.

25. Sturm, G., Finotello, F., Petitprez, F., Zhang, J.D., Baumbach, J., Fridman, W.H., List, M. and Aneichyk, T. (2019) Comprehensive evaluation of transcriptome-based cell-type quantification methods for immuno-oncology. Bioinformatics, 35, i436–i445.

26. Schelker, M., Feau, S., Du, J., Ranu, N., Klipp, E., MacBeath, G., Schoeberl, B. and Raue, A. (2017) Estimation of immune cell content in tumour tissue using single-cell RNA-seq data. Nature communications, 8, 1–12.

27. Newman, A.M., Liu, C.L., Green, M.R., Gentles, A.J., Feng, W., Xu, Y., Hoang, C.D., Diehn, M. and Alizadeh, A.A. (2015) Robust enumeration of cell subsets from tissue expression profiles. Nature methods, 12, 453–457.

28. Wagner, F., Yan, Y. and Yanai, I. (2018) K-nearest neighbor smoothing for high-throughput single-cell RNA-Seq data. BioRxiv, 217737.

29. Hafemeister, C. and Satija, R. (2019) Normalization and variance stabilization of single-cell RNA-seq data using regularized negative binomial regression. Genome biology, 20, 1–15.

30. Chen, B., Khodadoust, M.S., Liu, C.L., Newman, A.M. and Alizadeh, A.A. (2018), Cancer systems biology. Springer, pp. 243–259.

31. Gong, T. and Szustakowski, J.D. (2013) DeconRNASeq: a statistical framework for deconvolution of heterogeneous tissue samples based on mRNA-Seq data. Bioinformatics, 29, 1083–1085.

32. Dong, R. and Yuan, G.-C. (2021) SpatialDWLS: accurate deconvolution of spatial transcriptomic data. Genome biology, 22, 1–10.

33. Kleshchevnikov, V., Shmatko, A., Dann, E., Aivazidis, A., King, H.W., Li, T., Elmentaite, R., Lomakin, A., Kedlian, V. and Gayoso, A. (2022) Cell2location maps fine-grained cell types in spatial transcriptomics. Nature biotechnology, 1–11.

34. Kather, J.N., Heij, L.R., Grabsch, H.I., Loeffler, C., Echle, A., Muti, H.S., Krause, J., Niehues, J.M., Sommer, K.A. and Bankhead, P. (2020) Pan-cancer image-based detection of clinically actionable genetic alterations. Nature Cancer, 1, 789–799.

35. Fu, Y., Jung, A.W., Torne, R.V., Gonzalez, S., Vöhringer, H., Shmatko, A., Yates, L.R., Jimenez-Linan, M., Moore, L. and Gerstung, M. (2020) Pan-cancer computational histopathology reveals mutations, tumor composition and prognosis. Nature Cancer, 1, 800–810.

36. Jochems, C. and Schlom, J. (2011) Tumor-infiltrating immune cells and prognosis: the potential link between conventional cancer therapy and immunity. Experimental biology and medicine, 236, 567–579.

37. Taube, J.M., Klein, A., Brahmer, J.R., Xu, H., Pan, X., Kim, J.H., Chen, L., Pardoll, D.M., Topalian, S.L. and Anders, R.A. (2014) Association of PD-1, PD-1 ligands, and other features of the tumor immune microenvironment with response to anti–PD-1 therapy. Clinical cancer research, 20, 5064–5074.

38. Karaayvaz, M., Cristea, S., Gillespie, S.M., Patel, A.P., Mylvaganam, R., Luo, C.C., Specht, M.C., Bernstein, B.E., Michor, F. and Ellisen, L.W. (2018) Unravelling subclonal heterogeneity and aggressive disease states in TNBC through single-cell RNA-seq. Nature communications, 9, 1–10.

