## Supplementary Tables and Figures for "Cell type identification in spatial transcriptomics data can be improved by leveraging cell-type-informative paired tissue images using a Bayesian probabilistic model"

| | Linear Regression | CIBERSORT | spatialDWLS | DeconRNAseq | Stereoscope | GIST-base |
|---|---|---|---|---|---|---|
| Cell type 1 | 0.35 | 0.42 | 0.16 | 0.34 | 0.47 | 0.53 |
| Cell type 2 | 0.25 | 0.53 | 0.07 | 0.29 | 0.39 | 0.67 |
| Cell type 3 | 0.95 | 0.98 | NA | 0.92 | 0.96 | 0.82 |
| Cell type 4 | -0.27 | 0.65 | 0.13 | -0.22 | 0.59 | 0.63 |
| Cell type 5 | 0.29 | 0.34 | 0.11 | 0.22 | 0.23 | 0.33 |
| Cell type 6 | 0.98 | 0.98 | 0.84 | 0.98 | 0.98 | 0.57 |
| Overall MAE | 0.2147 | 0.0795 | 0.19 | 0.1275 | 0.0872 | 0.0839 |

**Supplementary Table S1:** Spearman's correlations values of estimated cell type proportions vs known ground truth for different methods in the Splatter-based simulations. Correlations are shown for each simulated cell types (rows 1-6). The overall mean absolute error (MAE) for the predictions is provided in the last row. For correlation, higher values indicate better performance; for MAE lower values are better.

| | Linear Regression | CIBERSORT | spatialDWLS | DeconRNAseq | GIST-base |
|---|---|---|---|---|---|
| B cell | 0.876 | 0.508 | 0.61 | 0.605 | 0.856 |
| Dendritic cell | -0.135 | -0.324 | -0.12 | -0.666 | 0.0450 |
| Macrophage/Monocyte | 0.493 | 0.166 | 0.21 | 0.0509 | 0.854 |
| NK cell | 0.919 | 0.608 | 0.58 | 0.917 | 0.832 |
| T cell CD4+ | 0.290 | 0.249 | 0.44 | 0.249 | 0.704 |
| T cell CD4+ (non-reg.) | 0.366 | -0.0151 | 0.42 | 0.112 | 0.435 |
| T cell CD8+ | 0.900 | 0.866 | 0.42 | 0.803 | 0.493 |
| T cell regulatory (Tregs) | -0.194 | 0.439 | 0.14 | -0.378 | 0.653 |
| Overall MAE | 0.1369 | 0.0946 | 0.1039 | 0.1080 | 0.0643 |

**Supplementary Table S2:** Like Table 1, but for the 2$^{nd}$ simulation study based on mixtures of real single-cell RNA-seq data, as described in Strum *et al*.

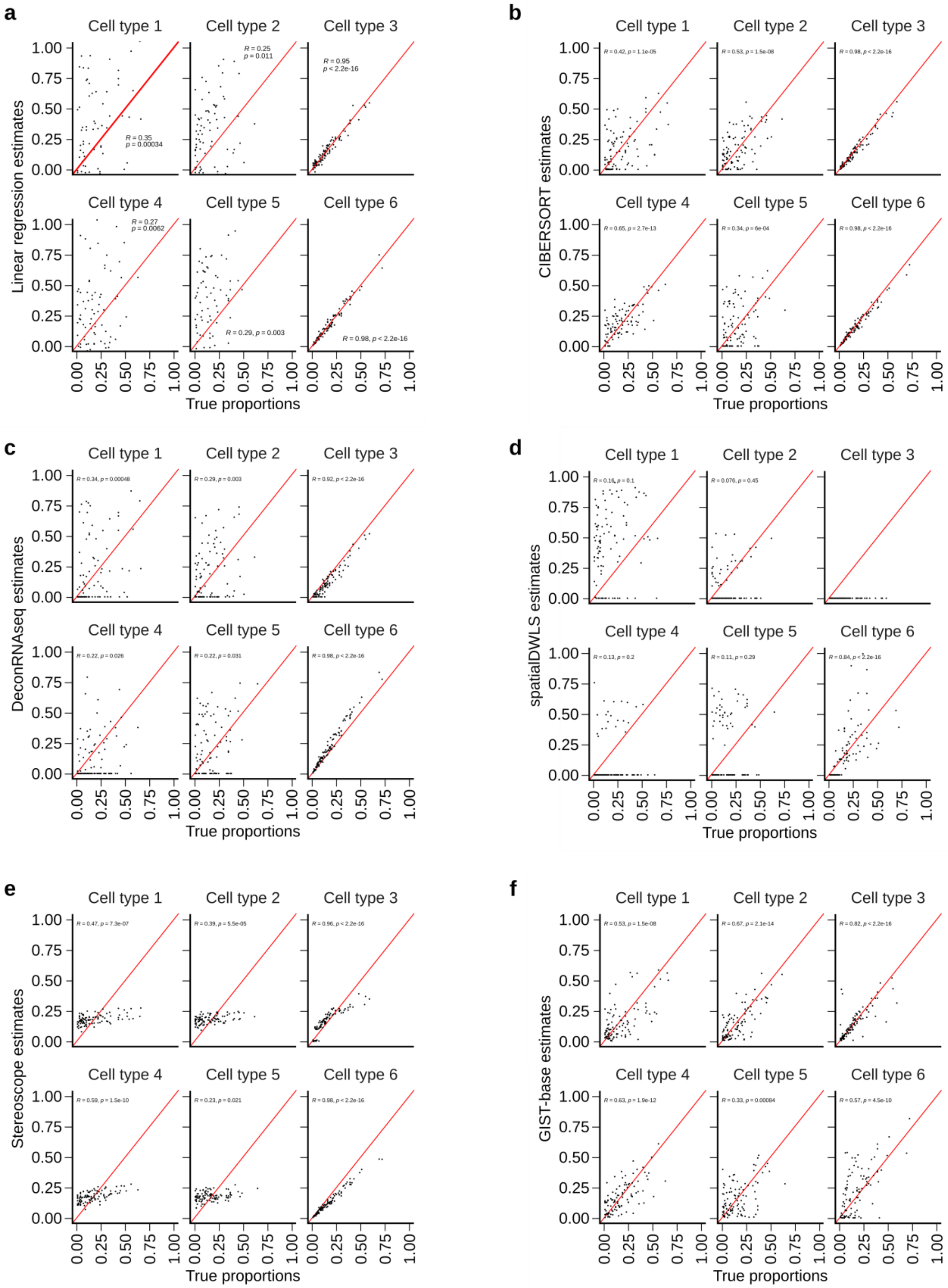

a
b
c
d
e
f
Cell type 1
Cell type 2
Cell type 3
Cell type 4
Cell type 5
Cell type 6
Linear regression estimates
CIBERSORT estimates
DeconRNAseq estimates
spatialDWLS estimates
Stereoscope estimates
GIST-base estimates
True proportions
0.00
0.25
0.50
0.75
1.00
R = 0.35, p = 0.00034
R = 0.25, p = 0.011
R = 0.95, p < 2.2e-16
R = 0.27, p = 0.0062
R = 0.29, p = 0.003
R = 0.98, p < 2.2e-16
R = 0.42, p = 1.1e-05
R = 0.53, p = 1.5e-08
R = 0.98, p < 2.2e-16
R = 0.65, p = 2.7e-13
R = 0.34, p = 6e-04
R = 0.98, p < 2.2e-16
R = 0.34, p = 0.00048
R = 0.29, p = 0.003
R = 0.92, p < 2.2e-16
R = 0.22, p = 0.026
R = 0.22, p = 0.031
R = 0.98, p < 2.2e-16
R = 0.16, p = 0.1
R = 0.076, p = 0.45
R = 0.13, p = 0.2
R = 0.11, p = 0.29
R = 0.84, p < 2.2e-16
R = 0.47, p = 7.3e-07
R = 0.39, p = 5.5e-05
R = 0.96, p < 2.2e-16
R = 0.59, p = 1.5e-10
R = 0.23, p = 0.021
R = 0.98, p < 2.2e-16
R = 0.53, p = 1.5e-08
R = 0.67, p = 2.1e-14
R = 0.82, p < 2.2e-16
R = 0.63, p = 1.9e-12
R = 0.33, p = 0.00084
R = 0.57, p = 4.5e-10

**Figure S1: Comparison of methods on the Splatter simulation dataset.**

a) Scatter plot of known ground truth proportions (x-axis) and linear regression predictions (y-axis) for the Splatter simulation.
b) Like (a) but for CIBERSORT.
c) Like (a) but for DeconRNAseq.
d) Like (a) but for spatialDWLS
e) Like (a) but for Stereoscope.
f) Like (a) bur for GIST-base model.

In all panels $R$ indicates the Spearman's correlation coefficient and the red line is the identity line.

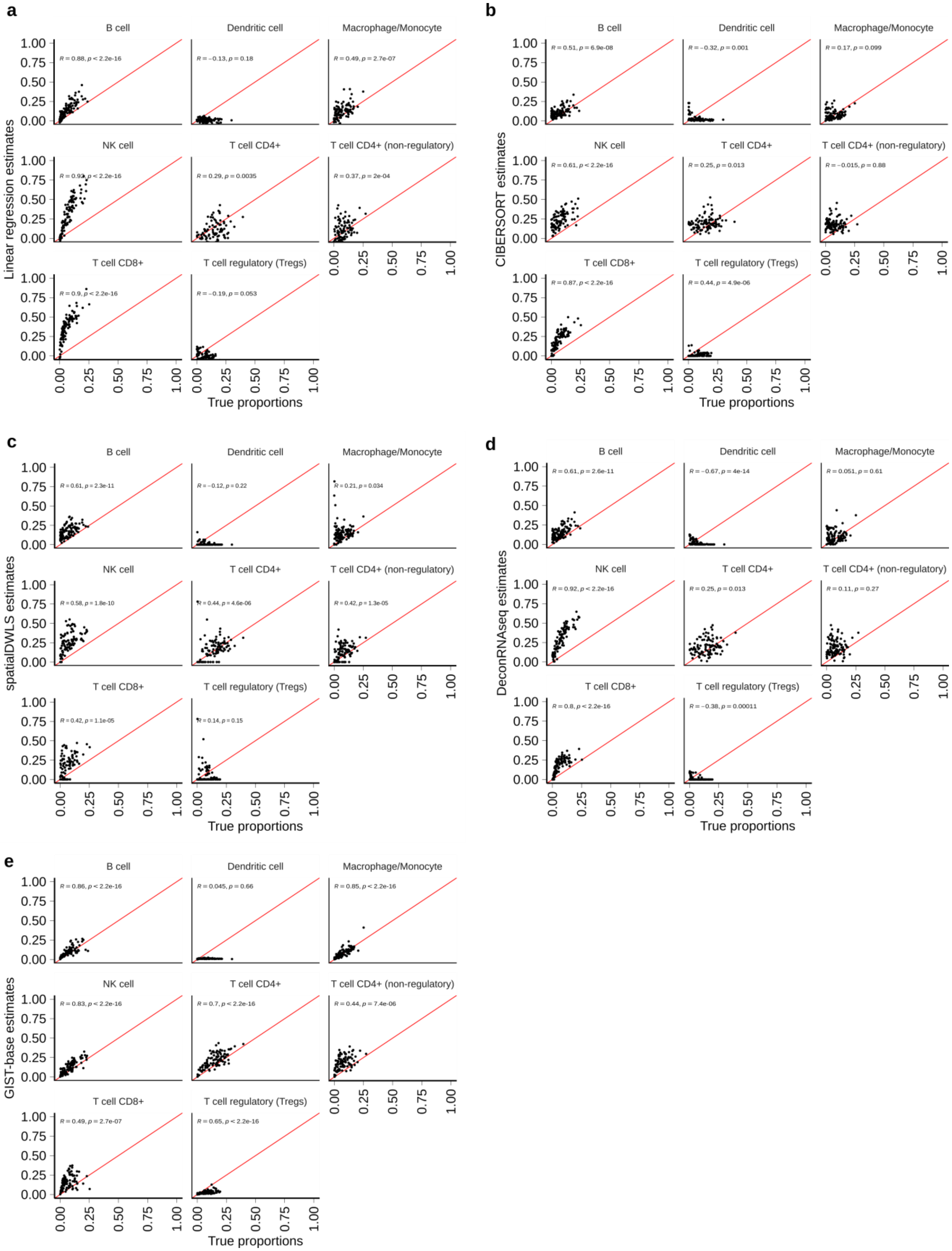

a
b
c
d
e
B cell
Dendritic cell
Macrophage/Monocyte
NK cell
T cell CD4+
T cell CD4+ (non-regulatory)
T cell CD8+
T cell regulatory (Tregs)
Linear regression estimates
CIBERSORT estimates
spatialDWLS estimates
DeconRNAseq estimates
GIST-base estimates
True proportions
0.00
0.25
0.50
0.75
1.00
R = 0.88, p < 2.2e-16
R = −0.13, p = 0.18
R = 0.49, p = 2.7e-07
R = 0.29, p = 0.0035
R = 0.37, p = 2e-04
R = 0.9, p < 2.2e-16
R = −0.19, p = 0.053
R = 0.51, p = 6.9e-08
R = −0.32, p = 0.001
R = 0.17, p = 0.099
R = 0.61, p < 2.2e-16
R = 0.25, p = 0.013
R = −0.015, p = 0.88
R = 0.87, p < 2.2e-16
R = 0.44, p = 4.9e-06
R = 0.61, p = 2.3e-11
R = −0.12, p = 0.22
R = 0.21, p = 0.034
R = 0.58, p = 1.8e-10
R = 0.44, p = 4.6e-06
R = 0.42, p = 1.3e-05
R = 0.42, p = 1.1e-05
R = 0.14, p = 0.15
R = 0.61, p = 2.6e-11
R = −0.67, p = 4e-14
R = 0.051, p = 0.61
R = 0.92, p < 2.2e-16
R = 0.25, p = 0.013
R = 0.11, p = 0.27
R = 0.8, p < 2.2e-16
R = −0.38, p = 0.00011
R = 0.86, p < 2.2e-16
R = 0.045, p = 0.66
R = 0.85, p < 2.2e-16
R = 0.83, p < 2.2e-16
R = 0.7, p < 2.2e-16
R = 0.44, p = 7.4e-06
R = 0.49, p = 2.7e-07
R = 0.65, p < 2.2e-16

**Figure S2: Comparison of methods on the Strum et al simulation dataset from mixtures of real single cell RNA-seq data.**

a) Scatter plot of known ground truth proportions (x-axis) and linear regression predictions (y-axis) for the Splatter simulation.
b) Like (a) but for CIBERSORT.
c) Like (a) but for spatialDWLS.
d) Like (a) but for DeconRNAseq.
e) Like (a) but for GIST-base model.

In all panels $R$ indicates the Spearman's correlation coefficient and the red line is the identity line.

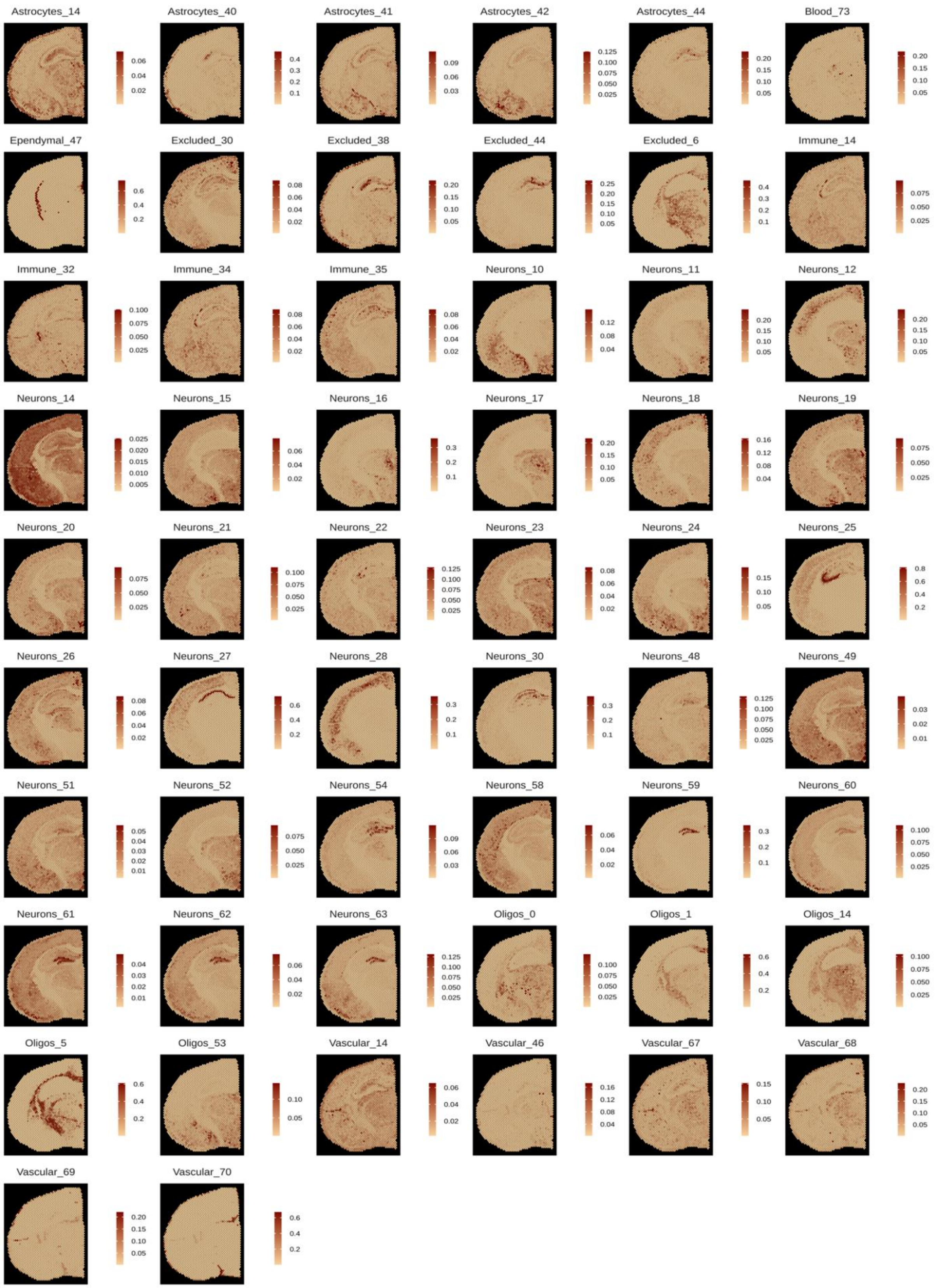


**Figure S3:** Proportion estimates for GIST base-model for 56 mouse brain cell subtypes. In the main text these cell subtypes were averaged to yield aggregate “glial” and “neuronal” estimates.

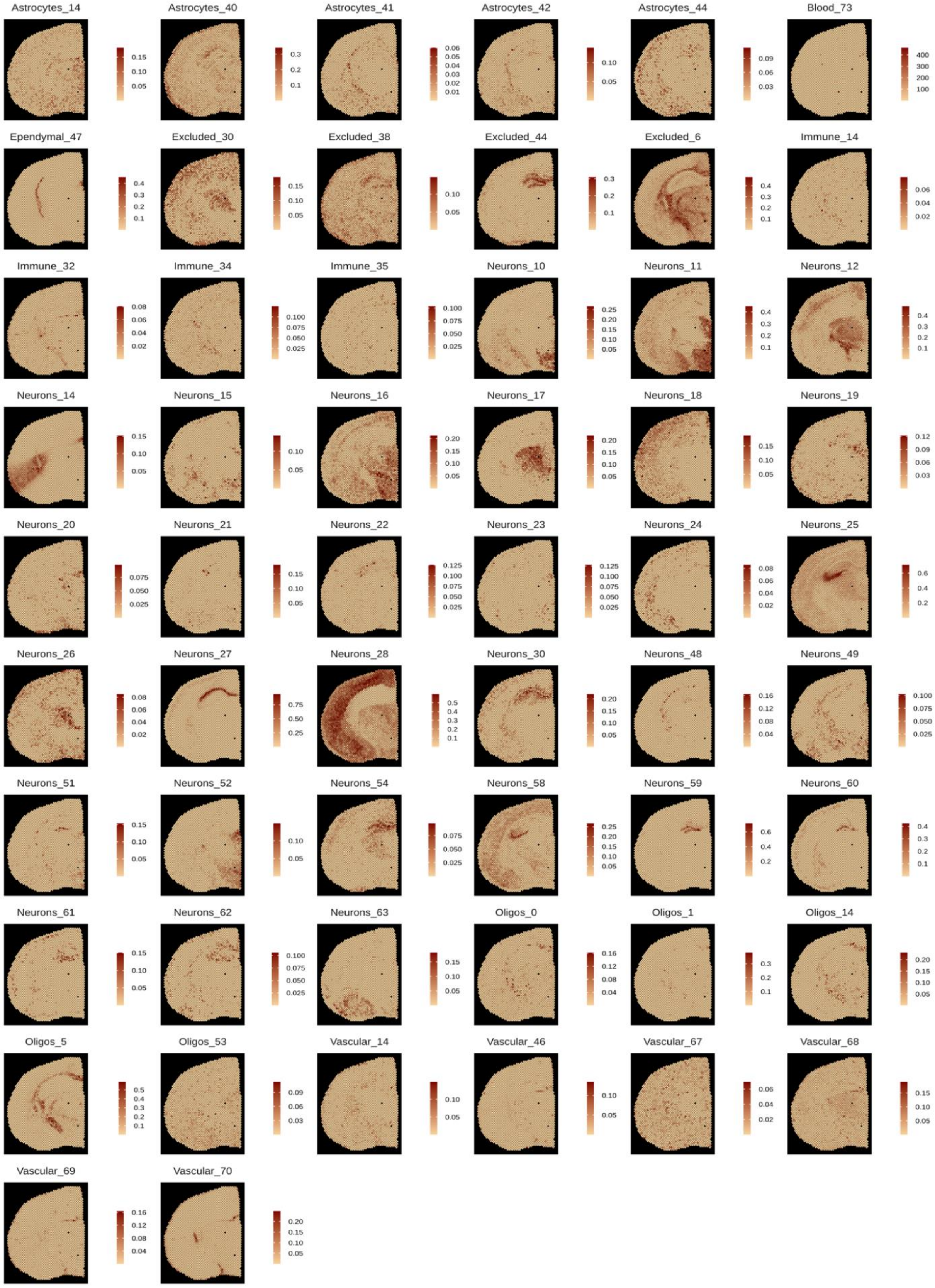

**Figure S4:** Like Fig. 3 but for the RCTD method.

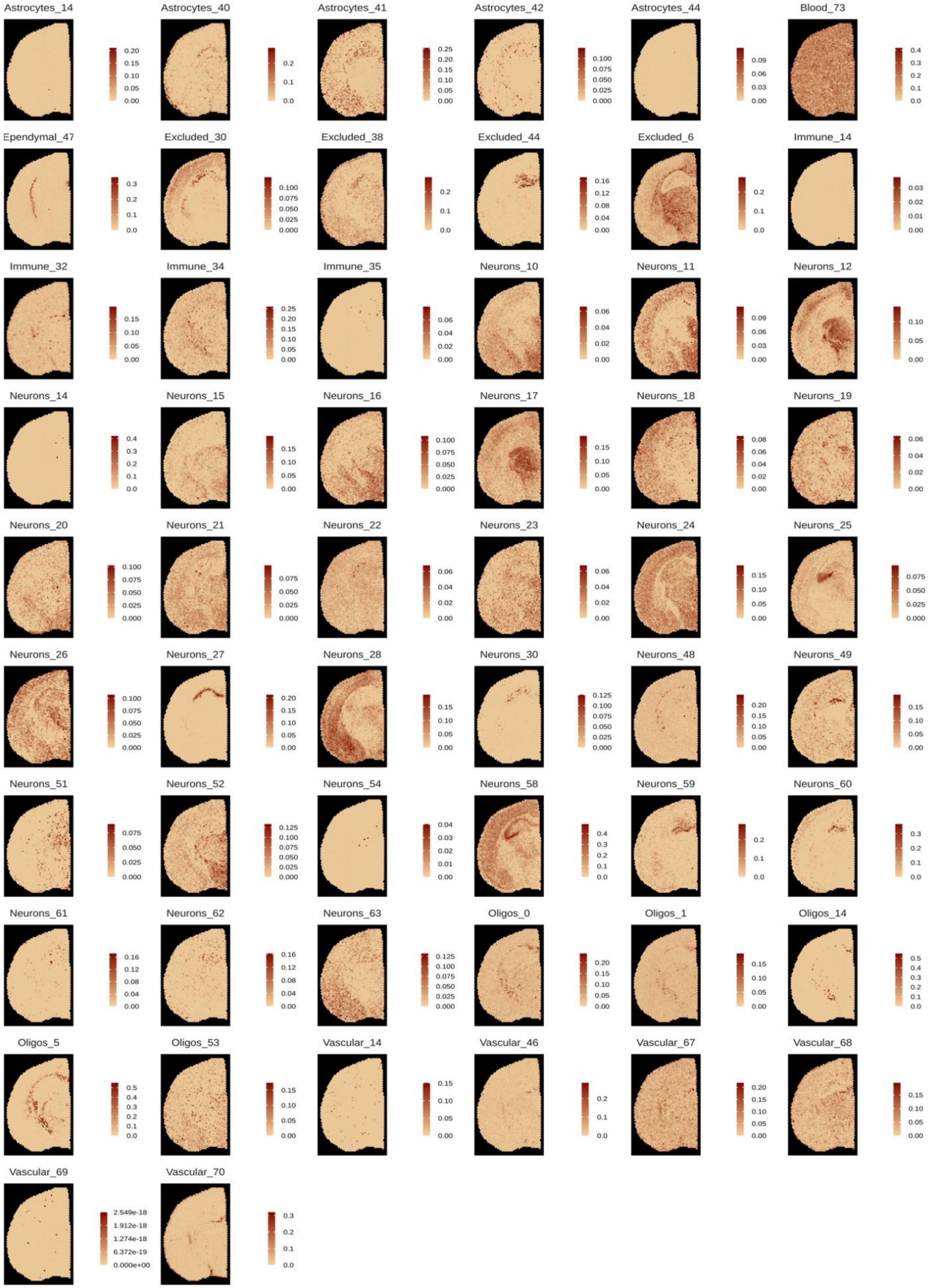


**Figure S5:** Like Fig. 3 but for the SPOTlight method.

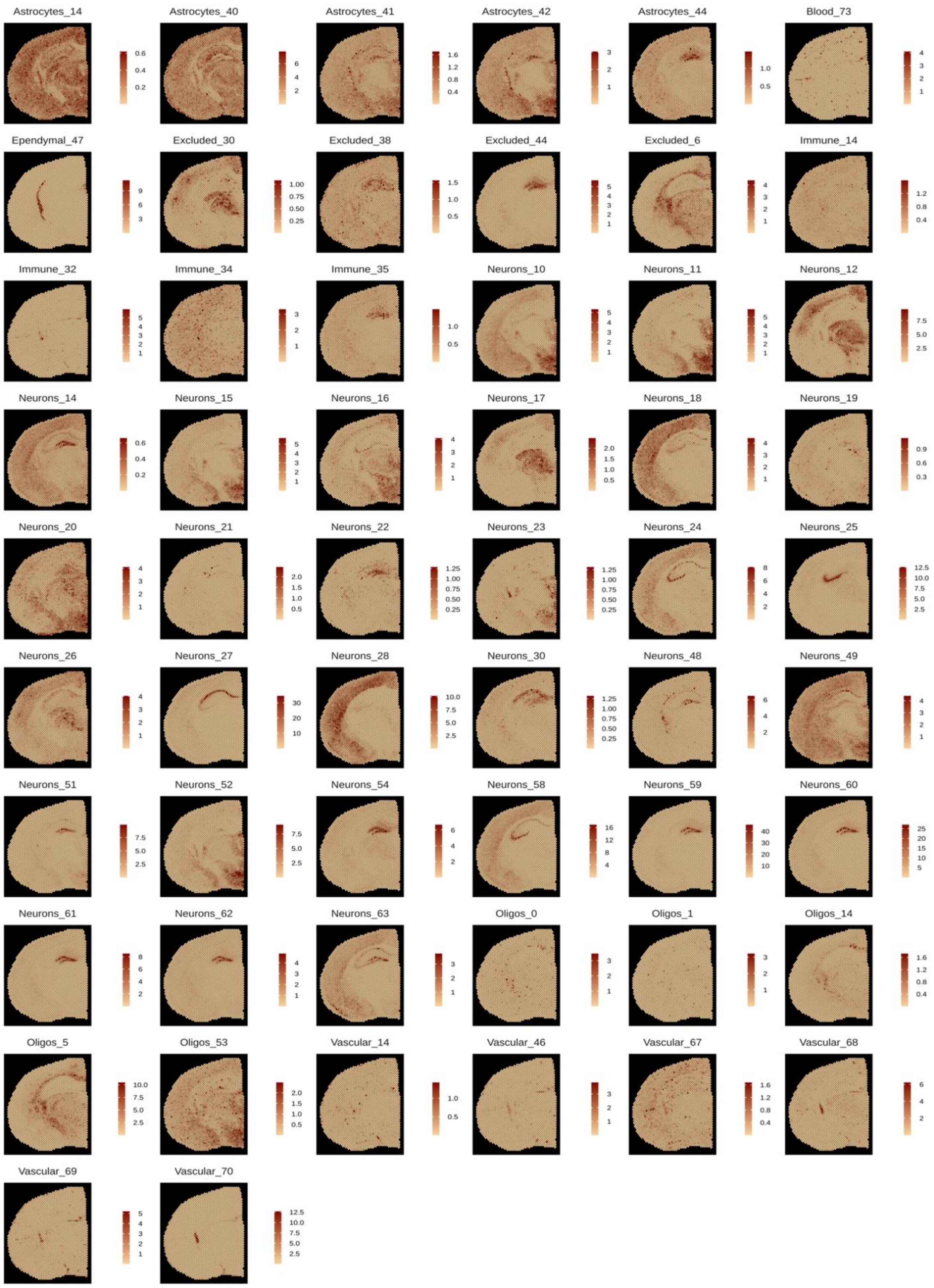


**Figure S6:** Like Fig. 3 but for the cell2location method.

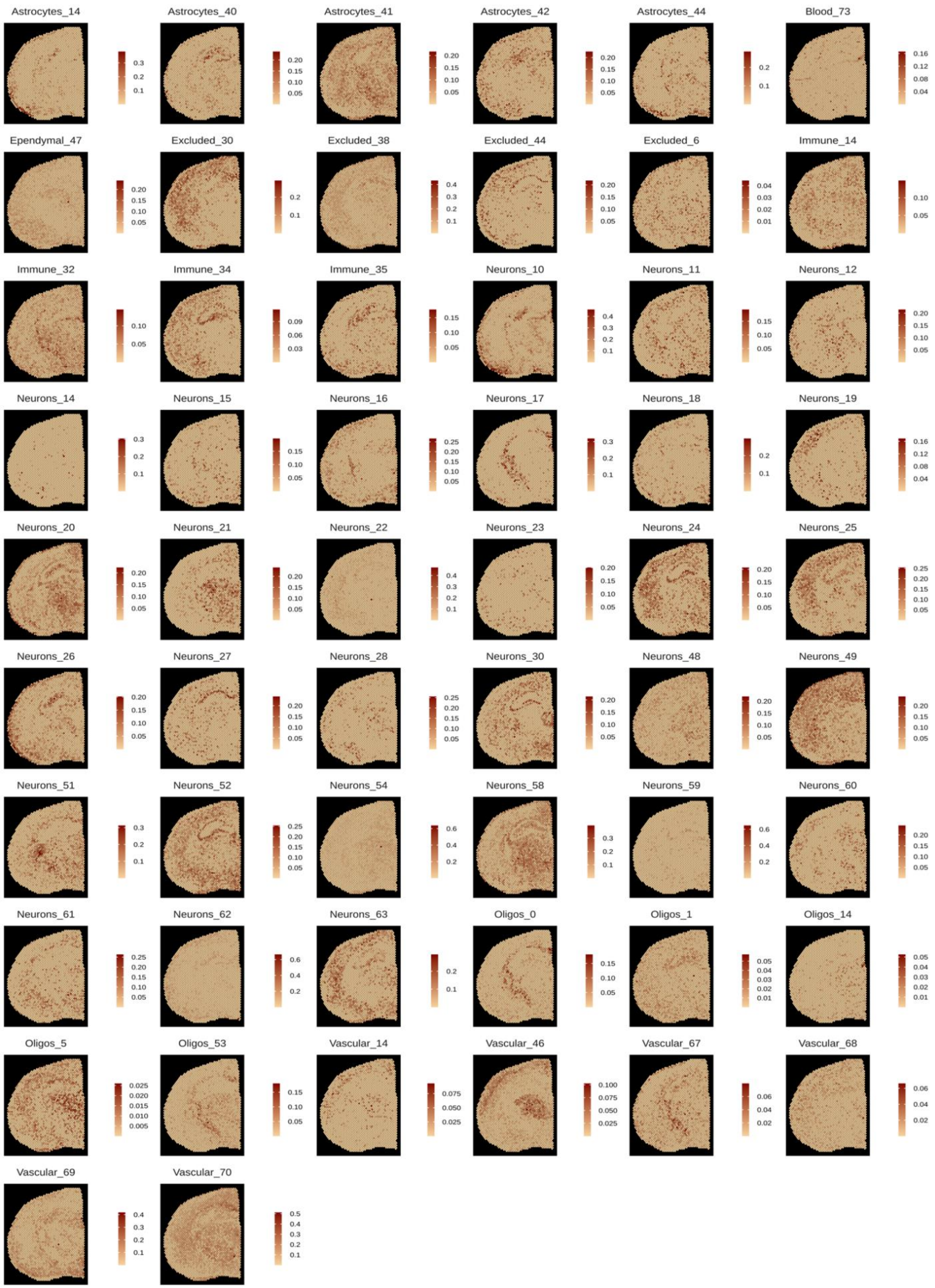


**Figure S7:** Like Fig. 3 but for the Stereoscope method.

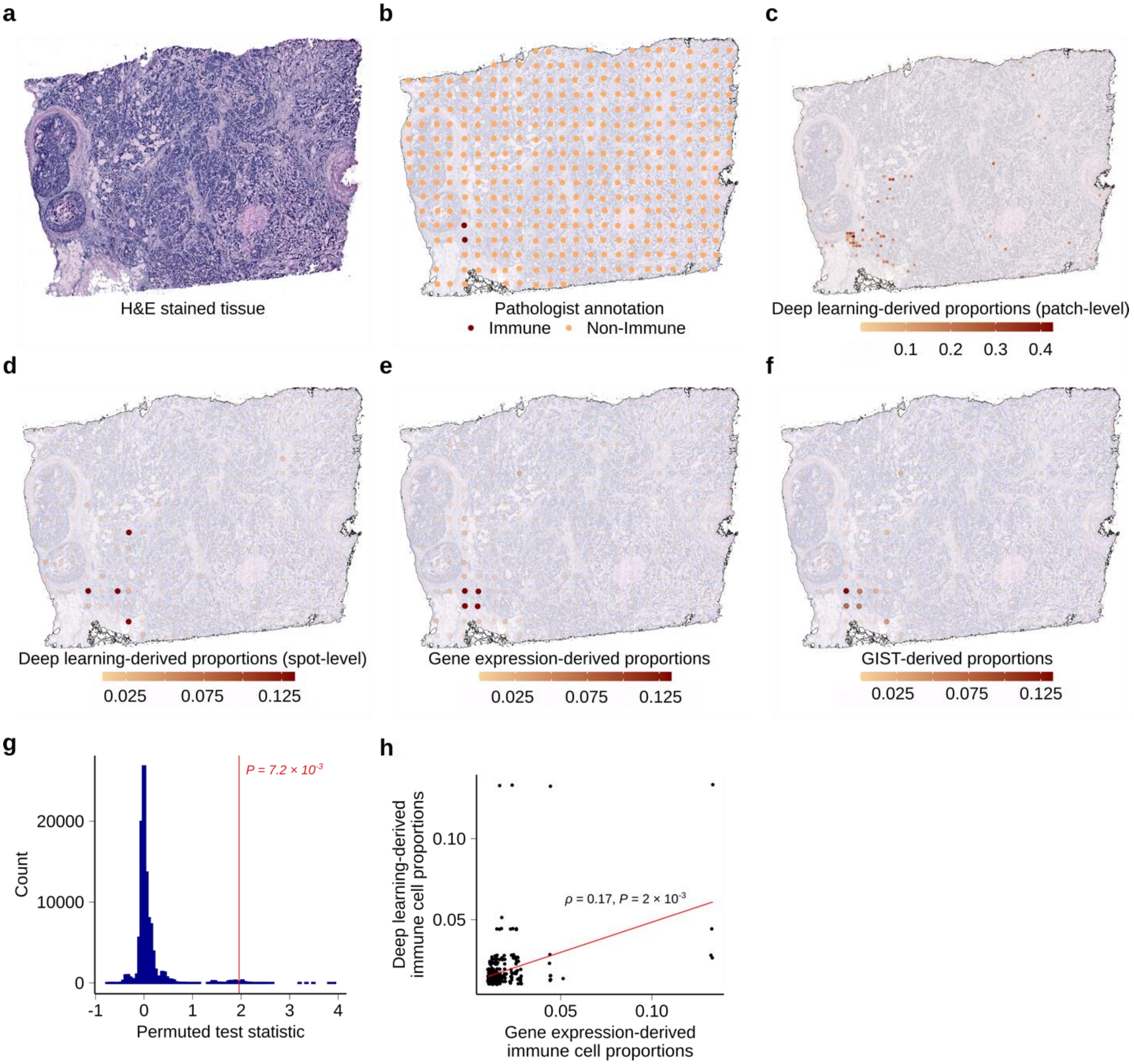


**Figure S8: Results summary for breast cancer Slide A1**

a) H&E stained tissue slide A1.
b) Pathologist annotation for slide A1 showing spots that were labelled immune cell infiltrated (marked by two dark colored spots). The image also shows all the spatial transcriptomics spots with their locations on the tissue.
c) Raw output from the deep learning model for slide A1 plotted on top of the tissue. The color scale shows immune cell predictions, made on 50 × 50 micron patches of the tissue.
d) Slide A1 with patch level deep learning predictions converted to spot level predictions. Spot level predictions are a sum of patch level predictions weighted by their overlap with the spot.
e) Slide A1 with only gene expression-derived immune cell proportions from the GIST base model.
f) Slide A1 with GIST-derived immune cell proportion relying on both expression and image-based modalities.

g) Histogram showing empirical null distribution of test statistic generated using a permutation procedure (x-axis, $\Delta_{perm}$). The test statistic is a measure of improvement by integration of image-based priors. The observed test statistic $\Delta$ is shown using a vertical red line.

h) Scatter plot showing correlation between deep learning-derived proportions (y-axis) and gene expression-derived proportions (x-axis). Each dot represents a spot on the tissue and red line is the regression line.

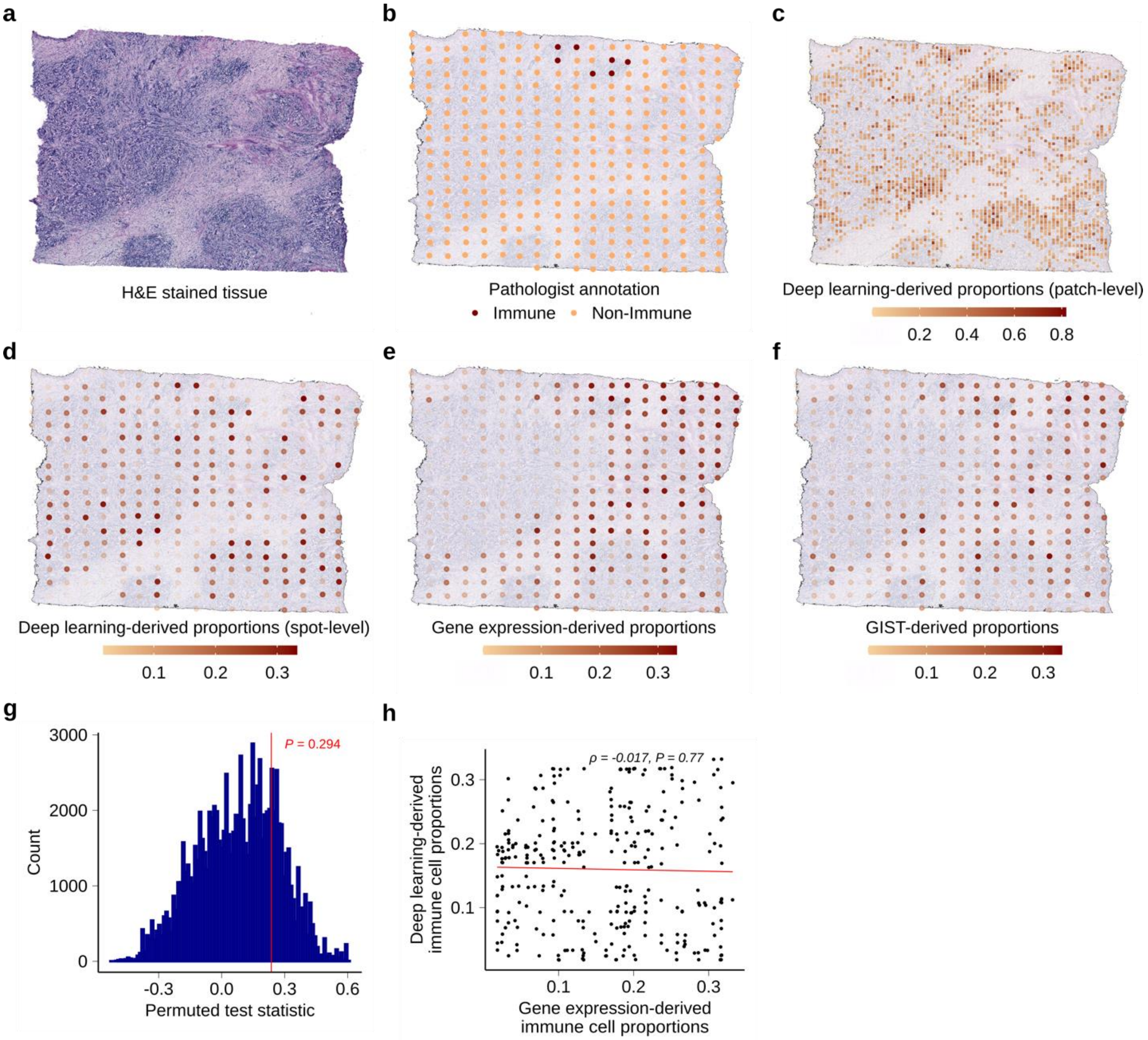


**Figure S9: Results summary for breast cancer Slide D1**

a) H&E stained tissue slide D1.
b) Pathologist annotation for slide D1 showing spots that were labelled immune cell infiltrated (marked by six dark colored spots). The image also shows all the spatial transcriptomics spots with their locations on the tissue.
c) Raw output from the deep learning model for slide D1 plotted on top of the tissue. The color scale shows immune cell predictions, made on 50 × 50 micron patches of the tissue.
d) Slide D1 with patch level deep learning predictions converted to spot level predictions. Spot level predictions are a sum of patch level predictions weighted by their overlap with the spot.
e) Slide D1 with only gene expression-derived immune cell proportions from the GIST base model.
f) Slide D1 with GIST-derived immune cell proportion relying on both expression and image-based modalities.

g) Histogram showing empirical null distribution of test statistic generated using a permutation procedure (x-axis, $\Delta_{perm}$). The test statistic is a measure of improvement by integration of image-based priors. The observed test statistic $\Delta$ is shown using a vertical red line.

h) Scatter plot showing correlation between deep learning-derived proportions (y-axis) and gene expression-derived proportions (x-axis). Each dot represents a spot on the tissue and red line is the regression line.

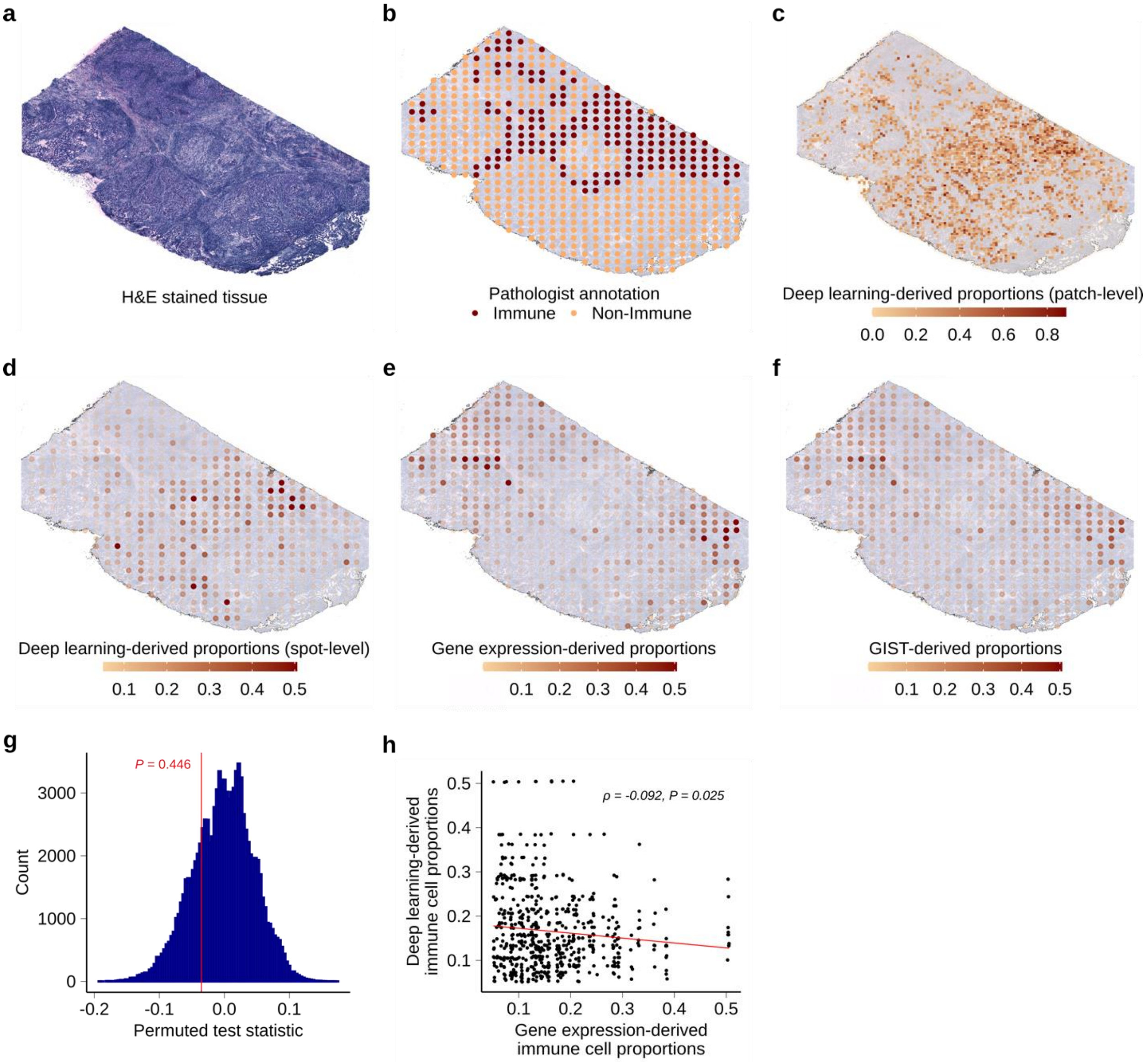


**Figure S10: Results summary for breast cancer Slide E1**

a) H&E stained tissue slide E1.
b) Pathologist annotation for slide E1 showing spots that were labelled immune cell infiltrated (marked by dark colored spots). The image also shows all the spatial transcriptomics spots with their locations on the tissue.
c) Raw output from the deep learning model for slide E1 plotted on top of the tissue. The color scale shows immune cell predictions, made on 50 × 50 micron patches of the tissue.
d) Slide E1 with patch level deep learning predictions converted to spot level predictions. Spot level predictions are a sum of patch level predictions weighted by their overlap with the spot.
e) Slide E1 with only gene expression-derived immune cell proportions from the GIST base model.
f) Slide E1 with GIST-derived immune cell proportion relying on both expression and image-based modalities.

g) Histogram showing empirical null distribution of test statistic generated using a permutation procedure (x-axis, $\Delta_{perm}$). The test statistic is a measure of improvement by integration of image-based priors. The observed test statistic $\Delta$ is shown using a vertical red line.

h) Scatter plot showing correlation between deep learning-derived proportions (y-axis) and gene expression-derived proportions (x-axis). Each dot represents a spot on the tissue and red line is the regression line.

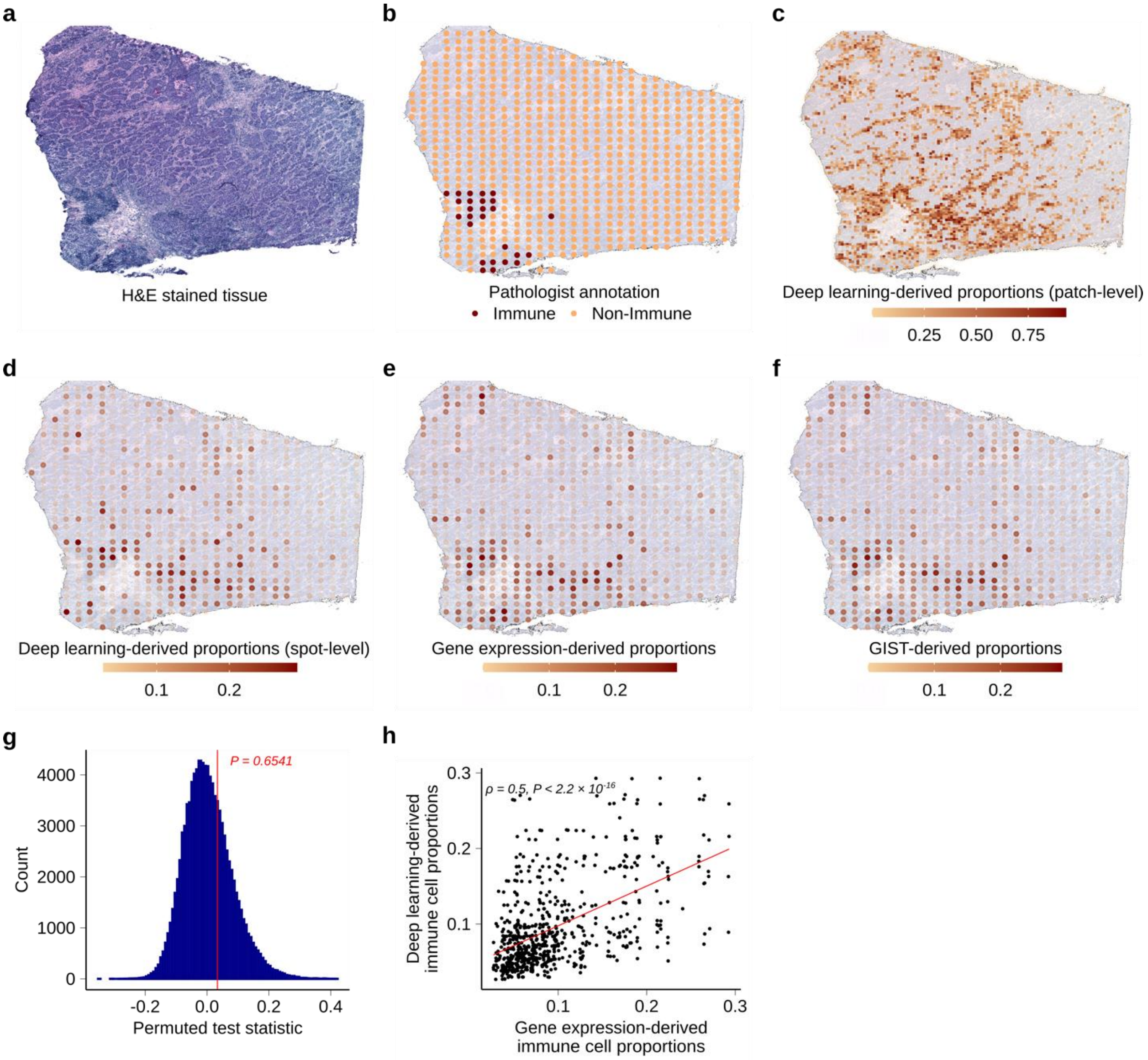


**Figure S11: Results summary for breast cancer Slide F1**

a) H&E stained tissue slide F1.
b) Pathologist annotation for slide F1 showing spots that were labelled immune cell infiltrated (marked by 26 dark colored spots). The image also shows all the spatial transcriptomics spots with their locations on the tissue.
c) Raw output from the deep learning model for slide F1 plotted on top of the tissue. The color scale shows immune cell predictions, made on 50 × 50 micron patches of the tissue.
d) Slide F1 with patch level deep learning predictions converted to spot level predictions. Spot level predictions are a sum of patch level predictions weighted by their overlap with the spot.
e) Slide F1 with only gene expression-derived immune cell proportions from the GIST base model.
f) Slide F1 with GIST-derived immune cell proportion relying on both expression and image-based modalities.

g) Histogram showing empirical null distribution of test statistic generated using a permutation procedure (x-axis, $\Delta_{perm}$). The test statistic is a measure of improvement by integration of image-based priors. The observed test statistic $\Delta$ is shown using a vertical red line.

h) Scatter plot showing correlation between deep learning-derived proportions (y-axis) and gene expression-derived proportions (x-axis). Each dot represents a spot on the tissue and red line is the regression line.

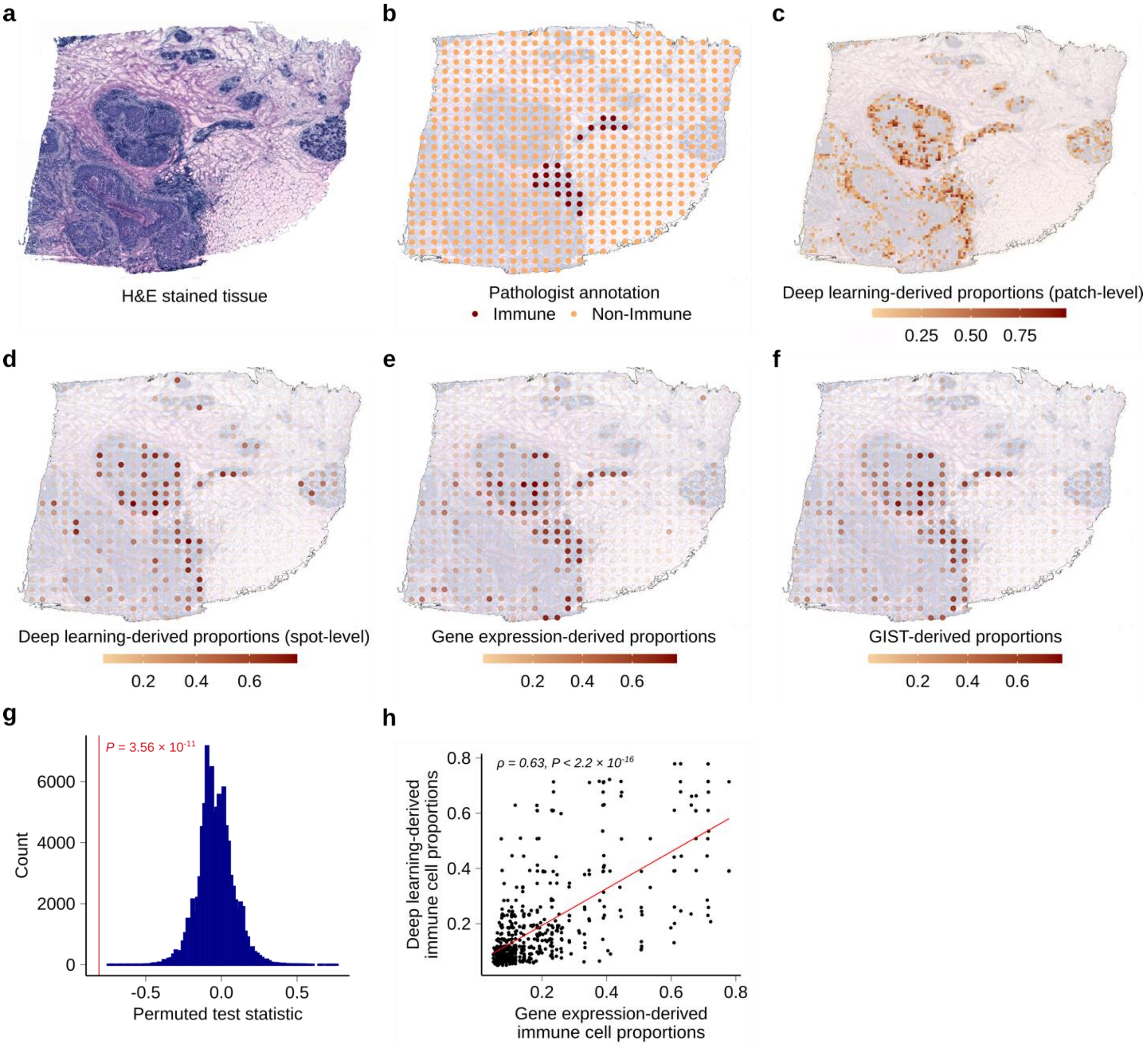


**Figure S12: Results summary for breast cancer Slide H1**

a) H&E stained tissue slide H1.
b) Pathologist annotation for slide H1 showing spots that were labelled immune cell infiltrated (marked by 23 dark colored spots). The image also shows all the spatial transcriptomics spots with their locations on the tissue.
c) Raw output from the deep learning model for slide H1 plotted on top of the tissue. The color scale shows immune cell predictions, made on 50 × 50 micron patches of the tissue.
d) Slide H1 with patch level deep learning predictions converted to spot level predictions. Spot level predictions are a sum of patch level predictions weighted by their overlap with the spot.
e) Slide H1 with only gene expression-derived immune cell proportions from the GIST base model.
f) Slide H1 with GIST-derived immune cell proportion relying on both expression and image-based modalities.

g) Histogram showing empirical null distribution of test statistic generated using a permutation procedure (x-axis, $\Delta_{perm}$). The test statistic is a measure of improvement by integration of image-based priors. The observed test statistic $\Delta$ is shown using a vertical red line.

h) Scatter plot showing correlation between deep learning-derived proportions (y-axis) and gene expression-derived proportions (x-axis). Each dot represents a spot on the tissue and red line is the regression line.

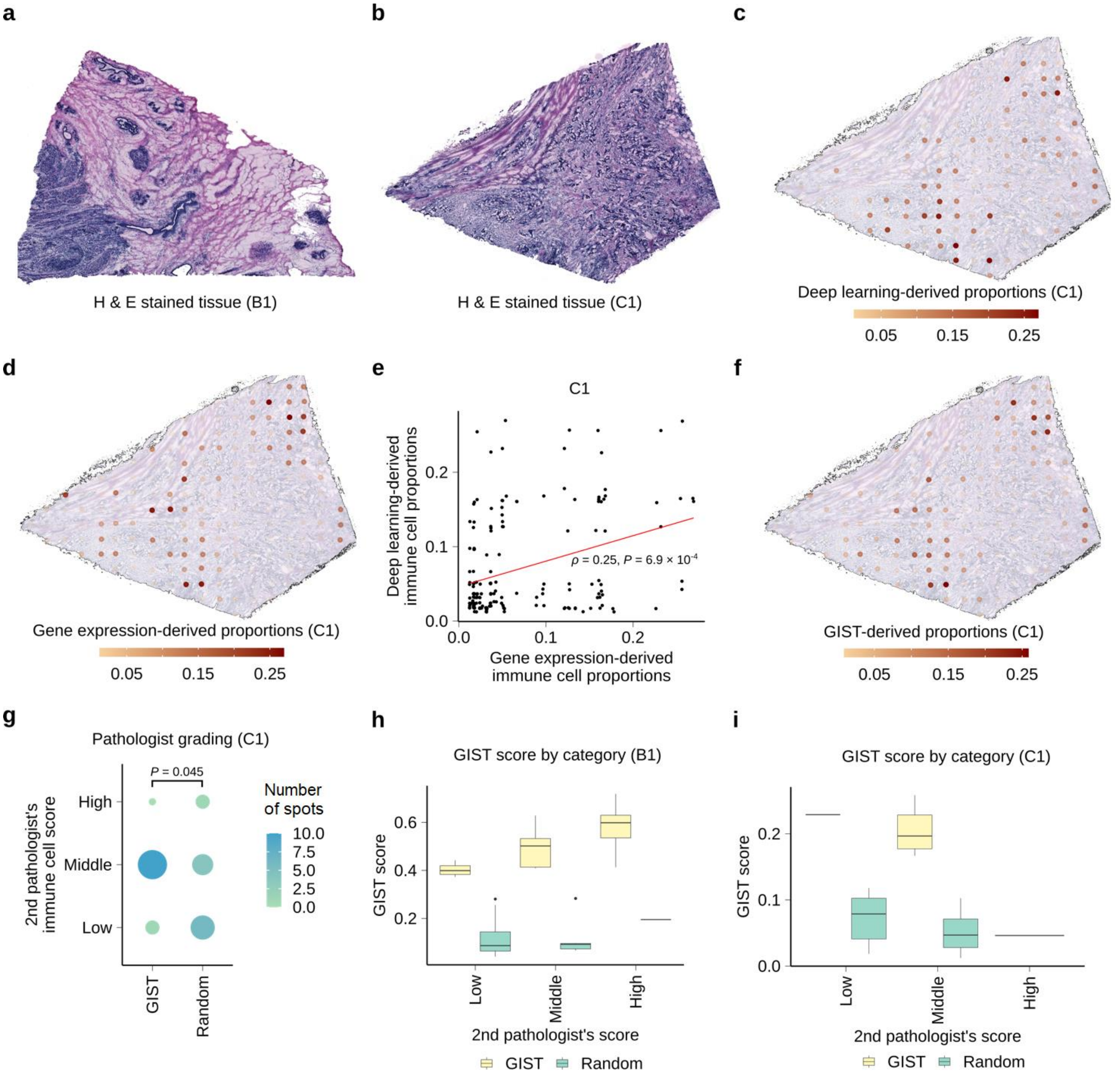


**Figure S13: 2nd Pathologist's reannotation of breast cancer slide's B1 and C1.**

a) H&E stained tissue for slide B1.
b) H&E stained tissue for slide C1.
c) Deep learning-derived proportions for slide C1. The color scale shows proportion of immune cells at a spot.
d) Spatial transcriptomics gene expression-derived proportions for slide C1. The color scale shows proportion of immune cells at a spot.
e) Scatter plot showing per-spot correlation between deep learning-derived predictions (y-axis) and gene expression-derived proportions (x-axis) for slide C1. Each dot is a spot and the red line is the regression line.
f) GIST-derived proportions for slide C1. The color scale shows proportion of immune cells at a spot.
g) Dot plot showing the second pathologist's immune infiltration grading with a score of low, middle and high (y-axis) for spots from different regions of the tissue (x-axis). Spots were

taken from slide C1 from high confidence regions from GIST model and random regions on the slide.

h) Boxplot showing distribution of GIST score (y-axis) broken down by immune infiltration grade (x-axis) provided by the second pathologist. For each pathologist grade (low, middle & high), GIST scores are shown for spots from annotated, GIST high confidence and random regions. Spots taken from slide B1.

i) Boxplot showing distribution of GIST score (y-axis) broken down by immune infiltration grade (x-axis) provided by the second pathologist. For each pathologist grade (low, middle & high), GIST scores are shown for spots from annotated, GIST high confidence and random regions. Spots taken from slide C1.
